# Recurrent mitotic error and purifying selection shape chromosomal mosaicism across human tissues

**DOI:** 10.64898/2026.10.27.760146

**Authors:** David Tang, Nolan Kamitaki, Yichen Si, Corinne E. Sexton, William C. Feng, Alexander D. Veit, Hye-Jung E. Chun, Peter J. Park, Po-Ru Loh

**Author notes:** Correspondence (D.T.), (P.-R.L.).

## Abstract

Mosaic chromosomal alterations (mCAs) have been extensively studied in cancer and clonal hematopoiesis but less so in healthy solid tissues. Here, we characterized mCAs in 19 tissues across 25 donors (aged 42–89+ years) by analyzing >200x whole-genome sequencing data generated by the Somatic Mosaicism across Human Tissues (SMaHT) Network. Across 722 tissue samples, we identified 1,008 autosomal mCAs >1 Mb in length and ubiquitous losses of sex chromosomes. Surprisingly, the highest levels of Y chromosome loss in males (5–10% cell fraction in elderly donors) were observed in the lung, adrenal gland, and heart. These tissues also tended to experience more X loss in females, revealing a gradient of mitotic error burden across human tissues. Autosomal aneuploidies displayed distinct tissue-specific patterns indicative of negative selection against most events, except in the adrenal gland, in which low-level aneuploidy was pervasive. In contrast to the adrenal gland, testis rarely harbored aneuploidies but frequently experienced low-level copy-neutral loss of heterozygosity (CN-LOH), suggesting increased mitotic recombination in spermatogonial stem cells. Recurrent X chromosome losses within tissues of female donors exhibited allelic skews explained by a combination of genetic and epigenetic effects. Together, these results show that the chromosomal landscape of aging human tissues is shaped by genetics and recurrent mutations, most but not all of which are held in check by purifying selection.

## Main

Cells in most human tissues acquire 10–50 point mutations in their genomes each year, such that most cells in adults contain hundreds to thousands of somatic point mutations^1,2^. Such mutations have been extensively studied, providing insights into developmental lineages^3,4^, mutagenic processes active in different tissues^5–7^, and mutational drivers of disease^8,9^. Mosaic chromosomal alterations (mCAs, also termed somatic copy number alterations) are another important but less-studied class of somatic mutations that produce megabase-to chromosome-scale changes in the chromosomal content of cells, via loss, gain, or copy-neutral loss of heterozygosity (CN-LOH). Although mCAs are much rarer than point mutations (and are not present in most cells), those that arise are much more likely to affect cellular function, survival, and proliferation. For this reason, chromosomal alterations and point mutations are the two major contributors to most cancers^10,11^. Aneuploidy in particular has generally been considered to be a characteristic of cancer, unusual in normal human tissues^12^.

Recent work has begun to change the notion that mCAs are primarily a feature of cancer genomes. Several lines of evidence have emerged indicating that chromosomal alterations in healthy tissues are much less rare than previously thought. In blood, which is readily accessible in population cohorts, SNP-array and bulk DNA sequencing studies have shown that mCAs (most frequently, sex chromosome losses) commonly affect a few percent of leukocytes of elderly individuals^13–18^. In healthy solid tissues, availability of tissue specimens is much more limited, but several key studies have begun resolving broadly similar, yet varying, mCA landscapes. Early studies observed a few aneuploid cells among ∼100 sequenced human neurons^19,20^ and hepatocytes^20^. Subsequently, a series of landmark studies sequenced millimeter-scale biopsies^21–23^, submillimeter microdissected specimens^4,6,7,24–29^, and single-cell-derived clonal cultures^5,23,30,3^ from one or more solid tissues of healthy donors. These studies observed recurrent losses of sex chromosomes^3,4^, common 9q CN-LOH mutations in skin^21^ and esophagus^22,23^, recurrent trisomies in gastric epithelium^29^, and other sporadic mCAs^3–7,21–30^. Most recently, two studies profiled aneuploidy across tens of thousands of breast epithelial cells, observing clonally expanded, recurrent aneuploidies in ∼3% of cells^31,32^.

While these studies are converging on a revised understanding that healthy tissues carry chromosomal alterations in a small percentage of cells, many questions remain. For most tissues other than breast, the sample sizes analyzed to date (typically hundreds of monoclonal populations) have been insufficient to discern generalizable patterns of mCAs, save for the most recurrent events. Additionally, varying methodological approaches (particularly relating to sex chromosome analyses) and cohort characteristics pose a challenge when comparing results across different tissues.

Bulk sequencing of biosamples collected across many tissues of post-mortem donors is a complementary approach that can mitigate the above challenges. Such an approach previously enabled a pan-tissue analysis of clonally expanded mCAs (>10% cell fraction) using RNA sequencing data from the GTEx project^33^. Here, we characterized mCAs at much greater sensitivity (minimum cell fraction 0.1%) across 19 tissue types of 25 donors by analyzing deep whole-genome sequencing (WGS) data generated by the Somatic Mosaicism across Human Tissues (SMaHT) Network^1^. This comprehensive, uniform analysis revealed tissue-specific patterns of abundant low-level mCAs throughout the human body, providing insights into the interplay between mutational and selective forces.

## Results

### Profiling mosaic chromosomal alterations across 1S tissue types of 25 donors

We analyzed bulk whole-genome sequencing data generated by the SMaHT Network for up to 19 tissues (blood, buccal, and 17 solid tissues; **Fig. 1a**) of 16 male and 9 female post-mortem donors (age at death 42–89+ years)^34^. Solid tissue samples from each donor were processed for sequencing by extracting 1.5-mm or 3-mm cores from frozen ∼1-cm-thick tissue aliquots (**Fig. 1b**). Two cores from each available solid tissue sample and two aliquots from each blood or buccal swab sample were independently sent for deep short-read WGS (median coverage ∼213x across 722 cores; for brevity, we refer to both solid and non-solid specimens as “cores” here and below). Long-read WGS data were also generated for most tissue samples (n=258 using Oxford Nanopore Technologies (ONT), median coverage 63x; n=237 using PacBio HiFi, median coverage 41x).

**Figure 1:**
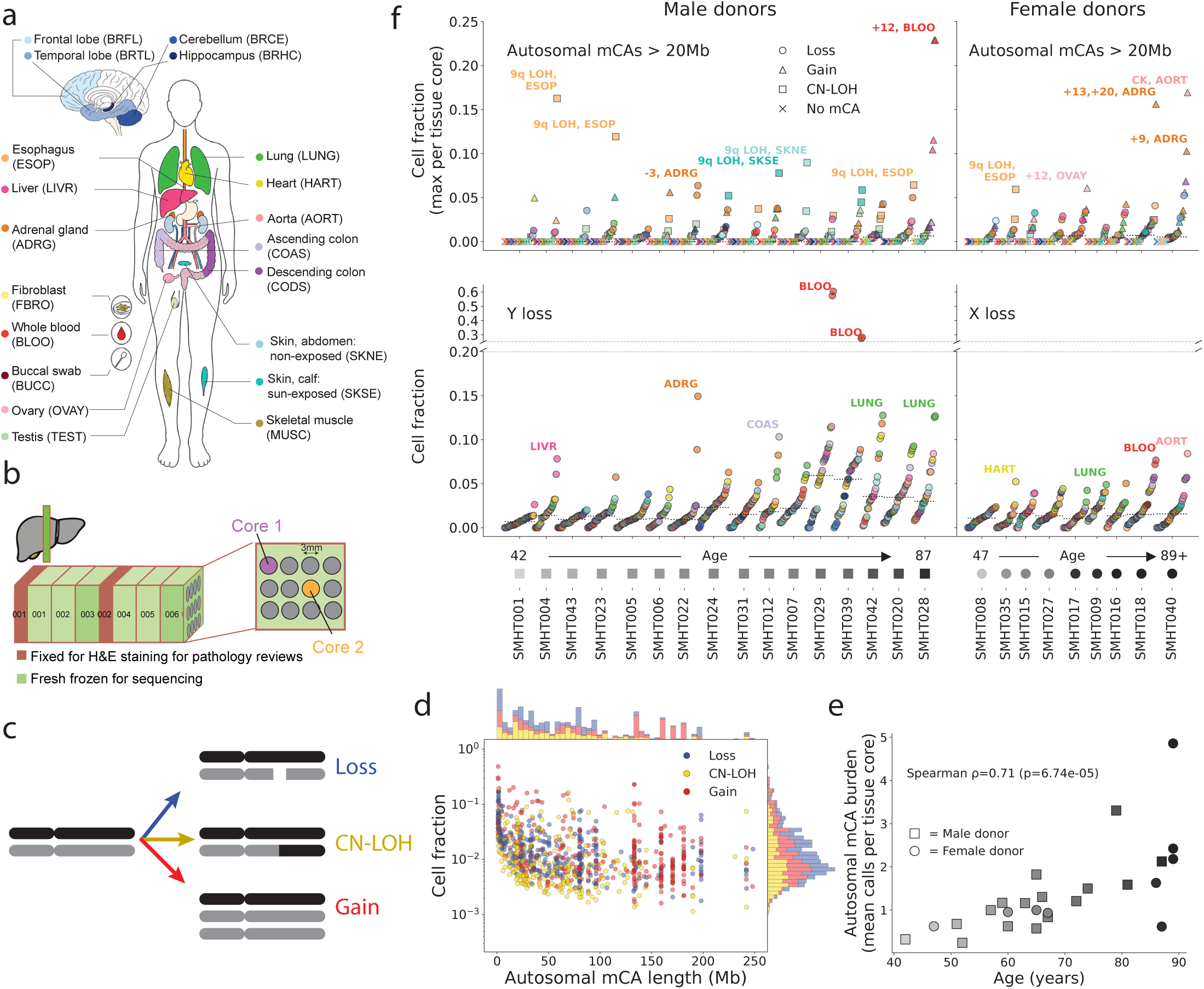
Body-wide landscape of mosaic chromosomal alterations in 25 donors. **(a)** Tissues sampled by SMaHT. Skin fibroblast cultures, which were sequenced at lower coverage, were excluded from mCA analysis, leaving 19 tissues (after merging left/right subcategories into bilateral categories for hippocampus, adrenal gland, ovary, and testis). **(b)** Example tissue coring protocol for solid tissues. Tissue cores of 1.5-mm or 3-mm diameter were extracted from flash-frozen tissue aliquots, typically of 1-cm thickness. For most tissues, two such cores were sent to two different sequencing centers for high-coverage (∼200x) short-read WGS. **(c)** Types of mCAs called by analyzing allelic imbalance and read depth in short-read WGS data. **(d)** Joint distribution of cell fraction and length of autosomal mCA calls. The lower envelope of the distribution reflects the limit of detection sensitivity, which improves with mCA length. CN-LOH mutations are detectable at lower cell fractions compared to gains and losses because CN-LOH generates twice the amount of allelic imbalance. Modes of the mCA length distribution correspond to whole-chromosome gains and losses. **(e)** Donor-level average number of autosomal mCAs per tissue core (across all cores available from a donor) versus donor age. **(f)** Donor-level summaries of large autosomal mCAs and sex chromosome losses found across tissues. Data are grouped by donor, with donors sorted left-to-right by sex followed by age (indicated at the bottom). Top: autosomal mCA status of each of a donor’s tissue cores. For each core in which at least one large (>20 Mb) autosomal mCA was detected, the highest cell fraction among these mCAs is plotted (with the copy-number state of the corresponding mCA indicated by marker shape: loss, circle; gain, triangle; CN-LOH, square); ‘X’ markers indicate cores with no large autosomal mCA. Data points in each donor series are sorted by cell fraction. Bottom: estimated LOY and LOX cell fractions in each tissue core from male and female donors, respectively. Images in panels **a** and **b** were adapted from ref. ^34^.

We adapted computational techniques that we recently developed and applied to blood WGS data^16,35^ to detect mCAs present within each tissue core (loss, gain, and CN-LOH events; **Fig. 1c**). To identify autosomal mCAs, we searched for allelic imbalances between a donor’s two haplotypes (**Supplementary Fig. 1**)—which we accurately phased (1 switch error per ∼30 Mb; **Supplementary Fig. 2**) by combining statistical phasing with long-read phase information. We then classified events as loss, gain, or CN-LOH based on read-depth within the event boundaries identified from allelic imbalance. For quantifying cell fractions of loss of the Y chromosome (LOY) in males, we determined that analyzing allelic imbalance in PAR1 was more robust than analyzing read-depth (**Supplementary Fig. 3**), though the two estimates were nonetheless correlated (**Supplementary Fig. 4**). However, read-depth analysis was more robust for quantifying X chromosome loss (LOX) in females, as recurrent loss of the two homologous X chromosomes produces competing allelic imbalances in opposite directions^35^.

Across the 722 tissue cores (8 to 50 (median 44) cores per tissue from 4 to 25 donors), we called 1,008 autosomal mCAs of length >1 Mb (median 63 Mb) and observed pervasive LOY in males and LOX in females (**Fig. 1d-f**). Autosomal mCA calls had a median cell fraction of 1% and a lower limit of 0.1% (**Fig. 1d**); for a spatially localized mutation, 0.1% cell fraction within a SMaHT tissue core corresponds to a ∼0.3-to 0.5-mm spheroid clone. Most autosomal mCAs were observed in only one core per tissue (**Supplementary Fig. 5**). As expected, autosomal mCA burden increased with age (Spearman *P*=6.7 x 10^-5^; **Fig. 1e**, **Supplementary Figs. 6 and 7**), as did amounts of LOY and LOX (Spearman *P*=1.3 x 10^-5^ and Spearman *P*=0.07; **Fig. 1f**, **Supplementary Fig. 8**). Other properties of the call set also supported the robustness of the mCA calls: read-depth deviations in mCA regions were largely consistent with allelic imbalances (**Supplementary Figs. G** and **10**), and for whole-chromosome aneuploidies, read-depth measurements in p and q arms were concordant (**Supplementary Fig. 11**). The genomic distributions of mCAs also exhibited striking tissue specificity (explored in detail below), a pattern not expected from germline CNVs or technical artifacts. Altogether, multiple lines of evidence support the validity of the mCA calls (**Supplementary Note 1**).

The cell fraction distribution of autosomal mCAs differed greatly from those of LOY and LOX. Most tissue cores from elderly donors (aged > 80 years) contained at least one large (>20 Mb) autosomal mCA, typically at a cell fraction of ∼1% (**Fig. 1f**). Only 24 out of 722 tissue cores contained large autosomal mCAs with a higher cell fraction of >5%, most of which involved common clonal expansions of 9q CN-LOH mutations in skin or esophagus^21,22^, aneuploidies in adrenal gland^33^, or trisomy 12 in blood^13,14^ (**Fig. 1f**). In contrast, LOY (in males) and LOX (in females) were present in 1-10% of cells of most tissues of elderly donors (**Fig. 1f**), extending previous observations in blood^36–38,17,18^ and other tissues^3,4,31,39,40^. LOY and LOX cell fraction distributions were much less right-skewed than autosomal mCA fractions (**Fig. 1f**), hinting at recurrent losses of sex chromosomes that steadily accumulate in mitotic cells with age (rather than sporadic clonal expansions).

The large amount of mCA data available from the 25 SMaHT donors (1,008 autosomal mCAs and measurements of sex chromosome loss in 722 tissue cores) provided an opportunity to further evaluate this and other hypotheses about mCA formation and accumulation, and to resolve subtler patterns in the mCA landscape that were previously undetectable.

### Shared tissue-specific propensities to pervasive LOX and LOY

First, we compared LOX levels across tissues within each female donor and LOY levels within each male donor to understand how and why different tissues experience different amounts of sex chromosome loss (**Fig. 2a,b**). The relative ranking of tissues was similar across donors, and interestingly, between LOX and LOY: for example, brain tissues (when available; BRCE, BRFL, BRTL, BRHC) tended to exhibit the lowest levels of LOX and LOY, whereas lung was often near the top (**Fig. 2a,b**). To assess this quantitatively while controlling for incomplete sampling of tissues across donors of various ages, we estimated the expected amounts of LOX and LOY in each tissue at a fixed reference age (70 years) by regressing cell fraction on age. This approach was suitable for all tissues other than blood, which had two outliers (clonally expanded LOY in 28% and 60% of cells in two donors; **Fig. 1f**), so for blood, we used data from ∼70-year-old participants in the All of Us (AoU) cohort^41^ whom we previously profiled for LOX and LOY^35^.

**Figure 2:**
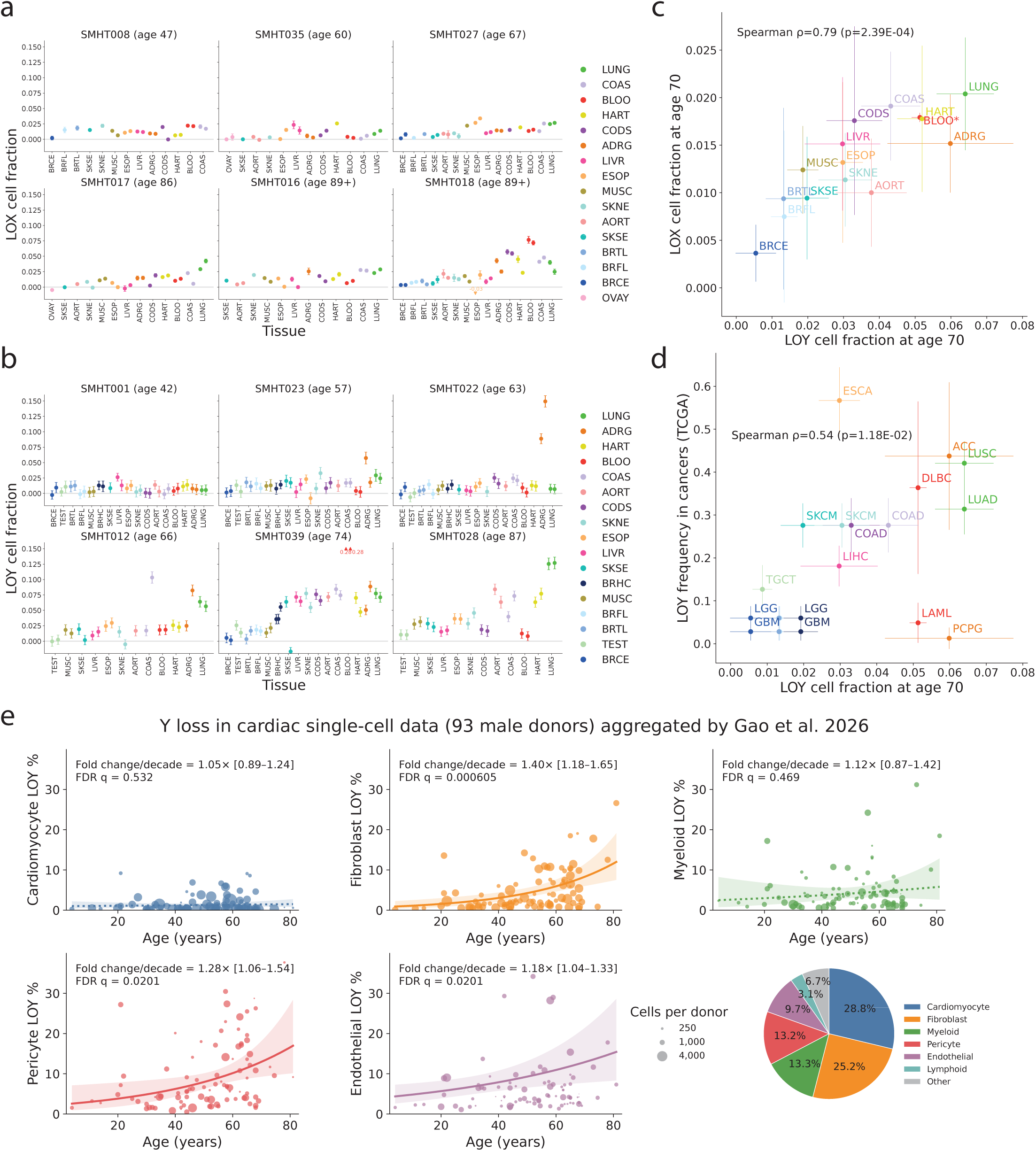
Loss of X and loss of Y are ubiquitous and elevated in the same tissues. **(a)** Cell fraction with loss of X in each available tissue core from 6 representative female donors, sorted by age. **(b)** Analogous data for loss of Y in male donors. In each panel, tissues are sorted (increasing left-to-right in plots and descending top-to-bottom in keys) in order of expected LOX (respectively, LOY) cell fraction at age 70 from linear regression of cell fraction on age (**Methods**). For LOX, we excluded tissue cores with imprecise estimates of LOX cell fraction (owing to high read-depth variability; median absolute deviation (MAD) > 0.05 among relative read-depth measurements in autosomal 1kb-bins), leaving only data from one core for some tissues. Additionally, analyses of LOX excluded hippocampus (BRHC) and analyses of both LOX and LOY excluded buccal samples (BUCC) due to insufficient sample size. **(c)** Age-adjusted X and Y loss frequencies (i.e., expected LOX and LOY cell fractions at age 70) plotted against each other. For blood, estimates are from ∼70-year-old participants in AoU^35^ rather than from regression of cell fraction on age in SMaHT. **(d)** Comparison of LOY cell fractions at age 70 in healthy tissues to rates of observing complete LOY in the dominant clone of primary tumors in TCGA^43^. Data are shown for tumor classes in TCGA with corresponding healthy tissues in SMaHT (**Supplementary Table 1**). Some tumors were mapped to multiple tissues (e.g., GBM to brain tissues, COAD to ascending and descending colon) and some tissues were mapped to multiple tumors (e.g., blood to DLBC and LAML, adrenal gland to ACC and PCPG). **(e)** Cell-type-specific LOY frequencies in left ventricle single-nucleus RNA-seq data from 93 adult male donors (in ref. ^46^) plotted against donor age. Each point corresponds to one cell type from a single donor with marker size scaling according to number of sequenced cells. The cell type composition of the atlas is broken down in the pie chart and only LOY frequencies in the most abundant cell types are plotted. The fitted line is from Poisson regression of LOY frequency on age, and the shaded area depicts its 95% confidence band. Error bars in all panels are 95% confidence intervals.

These analyses revealed that tissue-level propensities to LOX and LOY were strikingly correlated (Spearman *ρ*=0.79, *P*=2.4×10^-4^; **Fig. 2c**). Within each solid tissue, LOY levels (at age 70) were 2-to 4-fold higher than LOX levels, similar to blood^35^ (**Fig. 2c**). In blood, this difference appears to arise from a modest fitness-increasing effect of Y loss in males in contrast to fitness-neutral loss of the inactive^35,38,42^ X chromosome in females (leading to low-rate exponential growth of LOY versus linear accrual of LOX with age^35^). The SMaHT data suggest that this pattern may broadly hold across human tissues: i.e., in females, LOX steadily accumulates with age as cells occasionally experience mitotic error; and in males, the same error processes generate LOY, conferring a slight fitness advantage that is similar across tissues.

LOY levels in healthy tissues also correlated with frequencies of LOY in tumors arising from corresponding tissues (Spearman *ρ*=0.54, *P*=0.012; **Fig. 2d**). The role of Y chromosome loss in cancer remains unclear, but evidence is building that LOY may act as either a driver or a passenger event, with tendencies varying across tumor types^43,44^. The correlation with LOY levels in normal tissues could arise if genomic instability in cancer amplifies baseline levels of susceptibility to LOY. Additionally, the order-of-magnitude higher frequencies of LOY in most cancers relative to normal tissues (**Fig. 2d**) may point to LOY generally conferring a modest proliferative advantage that cancers readily harness.

The relative orderings of tissue-level propensities to LOX and LOY were somewhat unexpected (**Fig. 2a-c**, **Supplementary Figs. 12-14**). Previous work has suggested that LOY—a product of mitotic error—is mainly present in actively proliferating cells^45^. Consistent with this idea, blood and colon (tissues with high cell turnover rates) exhibited higher levels of LOY (and LOX), while the cerebellum (composed mostly of post-mitotic neurons) exhibited the least X and Y loss (**Fig. 2a-c**). However, the highest LOY levels were found in the lung, adrenal gland, and heart (which is generally thought to have low turnover), and these tissues also exhibited high levels of LOX (**Fig. 2c**). Moreover, the ranges in LOX and LOY levels across tissues were relatively narrow: only 2-fold for LOX and 3-fold for LOY across non-brain tissues (**Fig. 2c**). Given the much wider range in the lineage depths of different tissues (e.g., based on SBS1 mutations^2^), this suggests that rapidly dividing cell types have compensatory mechanisms that increase mitotic fidelity.

To validate and further explore the high levels of LOY in heart, we analyzed single-nucleus RNA sequencing data from the left ventricles of 93 adult male donors^46^. This allowed a probabilistic assessment of LOY within each cell based on its Y-chromosome gene expression. Similar to previous analyses of the brain^39^ and kidney^40^, LOY was enriched in specific cell types within the heart (fibroblasts, pericytes, and endothelial cells); in contrast, LOY fractions were minimal in cardiomyocytes and did not increase with age (**Fig. 2e**), consistent with cardiomyocytes being largely post-mitotic. Future single-cell data sets from other tissues will be needed to evaluate how cell type heterogeneity across tissues shapes measured levels of tissue-wide LOY, including the potential contribution of resident immune cells to bulk LOY measurements.

Finally, for a given tissue, levels of LOX or LOY within different tissue cores from the same donor tended to be more similar than between donors (**Fig. 2a,b**), even after adjusting for age (*P*=5.1 x 10^-4^ and *P*=1.2 x 10^-10^, respectively; **Methods**). This suggests that other donor characteristics—such as genetics^17,18,35^ and smoking^47,48^, which influence hematopoietic LOY—also influence LOX and LOY levels in solid tissues. Collectively, these results support a model in which each tissue of a donor has a predictable propensity to sex chromosome loss that is set by age, genetics, and lifestyle factors. Different tissue cores consisting of cells from different lineages nevertheless land close to this expectation owing to recurrent missegregation^4,49^.

### Tissue-specific patterns of common low-level autosomal aneuploidy

Autosomal mCAs provided distinct insights into mutation and selection in somatic cells. Most previous studies of autosomal mCAs in healthy solid tissues only had the sample size and sensitivity to observe common, clonally expanded mCAs^21–23,29^, with the broadest analysis to date finding 377 expanded mCAs (cell fraction ∼10% or higher)—dominated by 9q CN-LOH in esophagus and skin—in 2% of GTEx tissue samples (290 out of 16,672 samples)^33^. Here, the 1,008 autosomal mCAs (97% with cell fraction <10%; 54% with cell fraction <1%) that we identified in 722 deeply sequenced tissue cores revealed quite different, tissue-specific landscapes of autosomal mCAs commonly present at low cell fractions (**Fig. 3a,b**). Many specific events were observed several times among the 25 SMaHT donors, enough to identify patterns.

**Figure 3:**
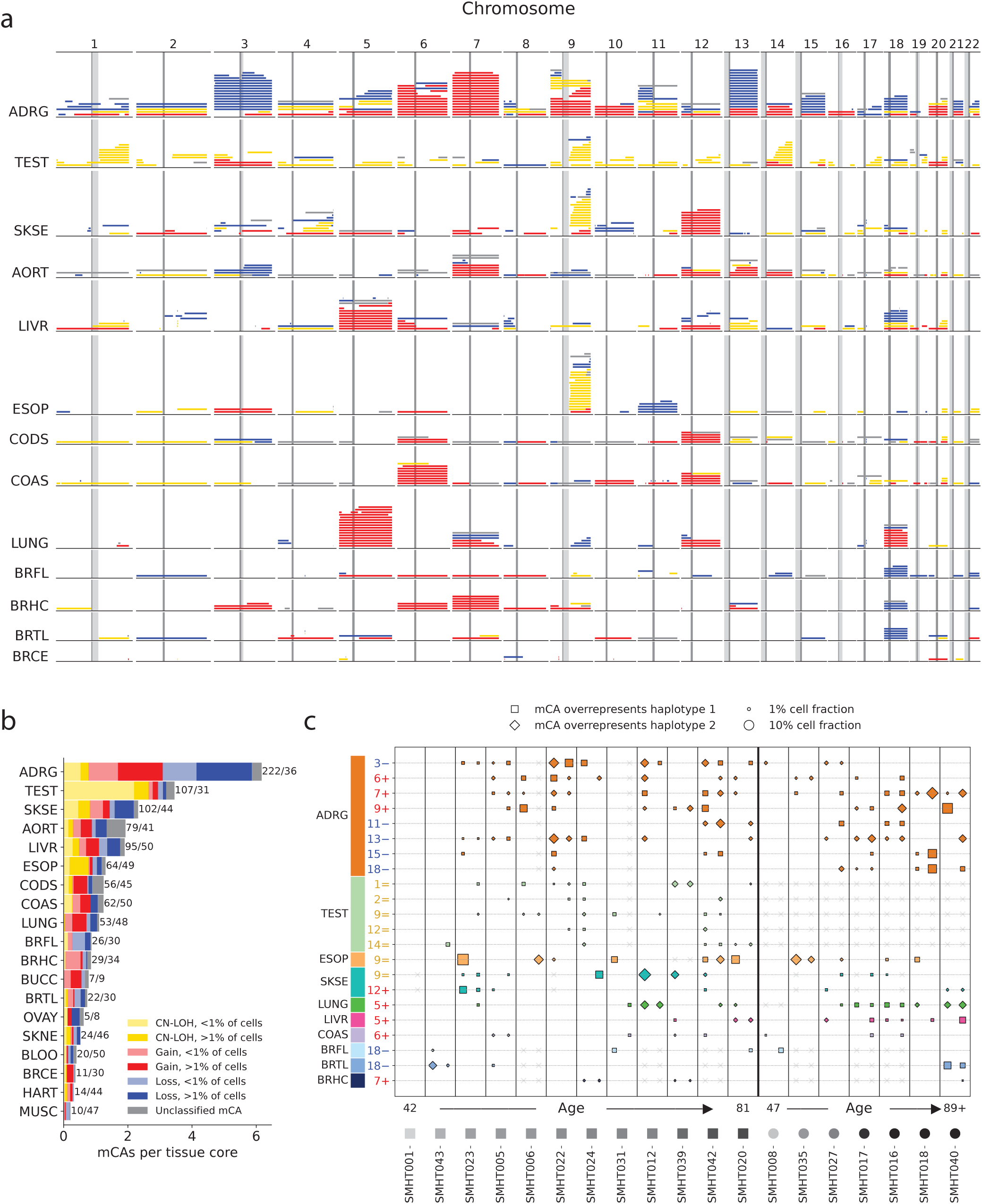
Tissue-specific patterns of autosomal mCAs. **(a)** Genomic locations of autosomal mCAs found in select tissues. Each horizontal line segment corresponds to a single mCA; mCAs detected on the same chromosome and in the same tissue of an individual are displayed at the same y-coordinate. Colors of line segments indicate copy-number state: loss, blue; CN-LOH, yellow; gain, red; unclassified, gray. **(b)** Average number of mCAs detected per core of each tissue type. Averages were computed as the number of detected mCAs divided by the total number of tissue cores (across all donors with the tissue available); these numbers are indicated to the right of each bar. Colors within the stacked bar plot indicate copy-number state, with mCAs of each class (CN-LOH, gain, loss) further stratified by cell fraction (lighter shading for cell fraction <1%; darker shading for cell fraction >1%). **(c)** Tissue-by-donor table of recurrent mCAs (classified by copy-number state and affected chromosome) present in >15% of cores from a tissue. Recurrent mCAs include several aneuploidies in adrenal gland, 1q, 9q, and 14q CN-LOH in testis, 9q CN-LOH in esophagus and skin, trisomy 5 in lung and liver, and loss of chromosome 18 in brain; copy-number states are abbreviated as –, =, and + for loss, CN-LOH, and gain, respectively. Pairs of columns present data for tissue cores from the same donor (typically two cores per tissue), with donors sorted left-to-right by sex followed by age (indicated at the bottom). Rows correspond to classes of recurrent mCAs, with markers within a row indicating the presence of an mCA of the indicated type in a specific core. Marker sizes scale with cell fraction and marker shapes (square or diamond) indicate the direction of allelic imbalance (toward haplotype 1 or haplotype 2) generated by the mCA. Gray ‘X’ markers indicate missing data (e.g., if a tissue was not present or not sampled from a donor). Donors without sequencing data from the adrenal gland were excluded from the plot. In the small fraction of instances in which three cores from a tissue were available from a donor, two of the three were randomly chosen to plot.

The most striking feature of the autosomal mCA call set was the amount of aneuploidy present in normal solid tissues (**Fig. 3a,b**, **Supplementary Fig. 15**). Most large mCAs (>20 Mb) were copy-altering (280 gains and 239 losses compared to 212 CN-LOH mutations), unlike in blood, in which most detectable mCAs are CN-LOH^16^. Overall 33% of tissue cores had at least one large mosaic gain or loss, with detection frequency varying across tissues (7% in cerebellum to 86% in adrenal gland). Aneuploidy was most dramatic in the adrenal gland, with most cores of older donors containing multiple aneuploidies each present in a few percent of cells (**Fig. 3c**). Low-level aneuploidy was also common in most other tissues (**Fig. 3a-c**), including brain tissues other than cerebellum (presumably reflecting aneuploid populations of non-neuronal cells, which comprise a much smaller fraction of the cerebellum^50^). These results generalize recent observations of recurrent aneuploidy in breast^31,32^ and gastric epithelium^29^, and suggest that most healthy human tissues of older individuals contain a small fraction (on the order of 1%) of aneuploid cells.

Aneuploidies in different tissues exhibited different genomic distributions, providing clues about the selective forces that shape aneuploidy in healthy tissues. In all tissues other than the adrenal gland, common aneuploidies (detected in multiple donors) were limited to at most a few chromosomes (**Fig. 3a**)—or none at all, such as in heart (**Supplementary Fig. 16**). This restriction on autosomal aneuploidies was observed despite recurrent mitotic missegregation in the heart and other tissues, as indicated by high rates of LOX and LOY (**Fig. 2c**), suggesting that unlike the Y chromosome and the inactive X chromosome, most autosomes are dosage-constrained in most tissues, with aneuploidies conferring enough of a fitness disadvantage for them not to be observed. Moreover, even the few common aneuploidies were typically only detected at low cell fractions, suggesting at most slight fitness advantages: although some aneuploidies were found in substantial fractions of tissue cores (e.g., trisomy 5 in lung and liver, trisomy 12 in skin, trisomy 6 in colon, loss of chromosome 18 in brain; **Fig. 3a**)—sometimes affecting each of an individual’s two haplotypes in different cores of the same tissue (**Fig. 3c**)—their cell fractions rarely exceeded 5% (**Fig. 3c**). Consistent with this observation, the autosomal aneuploidies observed in healthy tissues had limited overlap with cancers (**Supplementary Fig. 17**). Taken together, these results suggest that the overall landscape of aneuploidy in healthy tissues is driven by recurrent mitotic errors (probably involving all chromosomes), the products of which are then pruned by purifying selection that eliminates most autosomal aneuploidies. The adrenal gland appears to be a notable exception that is surprisingly tolerant of aneuploidy; conversely, purifying selection may be particularly efficient in blood (owing to its large effective population size^49^), potentially explaining its relatively lower rate of aneuploidy.

### Low-level accumulation of pre-meiotic crossovers in testis with age

CN-LOH mutations, which are generated by homologous recombination (HR) during DNA double-strand break (DSB) repair, displayed a very different mutational landscape: testis samples were surprisingly enriched for CN-LOH mutations throughout the genome (**Fig. 3a,b**). Testis tissue cores typically contained multiple detectable CN-LOH mutations (mean 2.9, range 0-12; **Fig. 3b,c**), increasing with age (Spearman *P*=0.02; **Supplementary Fig. 6**); in contrast, aneuploidy rates were lower in testis than most other tissues (**Fig. 3b**). All 82 of the CN-LOH mutations found in testis were present at low cell fractions (mean 0.7%, range 0.1-2.2%; **Fig. 3b,c**), indicating minimal effects on clonal fitness. Nonetheless, these events did appear clonal based on the presence of step-changes in allelic balance (**Supplementary Fig. 18**). This observation, together with the accrual of CN-LOH clones with age, suggests that these mCAs are unrelated to selection on sperm^51,52^ and instead reflect mitotic crossovers that occur in cycling spermatogonial stem cells. Consequently, spermatids may occasionally contain one or more crossovers of pre-meiotic origin. In contrast to pre-meiotic non-crossover gene conversions^53^, these pre-meiotic crossovers comprise a very small fraction of the ∼25 crossovers per human spermatid^54^, reflecting the strong bias toward DSB repair using non-crossover pathways in non-meiotic cells. The reason for the pronounced enrichment of mitotic crossovers in testis relative to other tissues remains a mystery; it is unclear whether this is related in some way to the upregulation of crossovers during meiosis later in the spermatogonial lineage.

The genomic distribution of CN-LOH mutations in testis also provided insight into the mutational forces underlying recurrent 9q CN-LOH in esophagus and skin. Assuming that CN-LOH mutations in testis experience minimal clonal selection, their breakpoint distribution reflects variation in propensity of different genomic regions to experience DSBs and subsequent HR-mediated repair. Consistent with this, the two hotspots of CN-LOH in testis—the late-replicating heterochromatic regions 1q12 and 9q12 (**Fig. 3a**)—are also hotspots of sister chromatid exchange (HR-mediated repair using the sister chromatid)^55^.

This suggests that the remarkable prevalence of 9q CN-LOH in esophagus and skin (as well as embryonic stem cells^56^)—which also show breakpoint enrichment in 9q12^33,57^—is driven not only by strong clonal selection on the *NOTCH1*-damaging mutations on which they act^21–23^, but also by the location of *NOTCH1* (near the 9q telomere): this locus may be among the most likely in the human genome to be affected by CN-LOH. Given the apparent anti-tumorigenic effect of *NOTCH1*-mutant clonal expansions^58^, this may be a fortunate coincidence.

### Remnants of early developmental chromosomal instability in adult tissues

While most somatic mutations in adults accrue over years of aging, a disproportionate fraction of somatic SNVs occur during early embryonic development and are propagated from progenitor cells to their descendants^3,4,59^. Chromosomal alterations are also common in early development but subsequently undergo strong filtering, both at the organismal level (as high levels of aneuploidy make embryos inviable) and the cellular level (from reduced clonal fitness)^60,61^. Early mCAs do sometimes propagate, as evidenced by mosaic cases of rare genetic disorders^59^, but the extent to which early mCAs are found in healthy adults is largely unexplored. Among the GTEx cohort profiled for high-cell-fraction mCAs, 1 donor out of 948 was found to have a body-wide CN-LOH of chromosome 10^33^.

Comprehensive WGS-based profiling of mCAs across tissues in SMaHT showed that persistence of early developmental mCAs at low cell fractions across adult tissues is much more common than previously appreciated. Three of the 25 donors had clear evidence (maximum cell fraction >2% with consistent breakpoints across tissues) of such mCAs: a 20q CN-LOH mutation, a deletion of 12p, and a 1.4-Mb *NF1* microdeletion (**Fig. 4**). A fourth donor had chromosome 18 loss detected at <1% cell fraction in several tissues (**Supplementary Fig. 1G**). The 20q CN-LOH was present in nearly all available tissues at modest cell fractions (<2% for most tissues; 3-4% in esophagus; **Fig. 4a**), while the 12p loss was observed at lower cell fractions in only about half of tissues, perhaps indicating a slight fitness cost (**Fig. 4b**). In contrast, the *NF1* deletion appeared to confer a proliferative advantage, as it was observed in every tissue at much higher cell fractions, up to 16% in blood (**Fig. 4c**). This did not reflect leukocyte presence in solid tissues, as the same donor also had an expanded hematopoietic trisomy 12 detected at much lower cell fractions in solid tissues (bounding the amounts of blood infiltration; **Fig. 4d**). This donor did not have a neurofibromatosis type 1 diagnosis, consistent with mosaic *NF1* deletion being a low-penetrance mutation observed in healthy population cohorts^16,62^.

**Figure 4:**
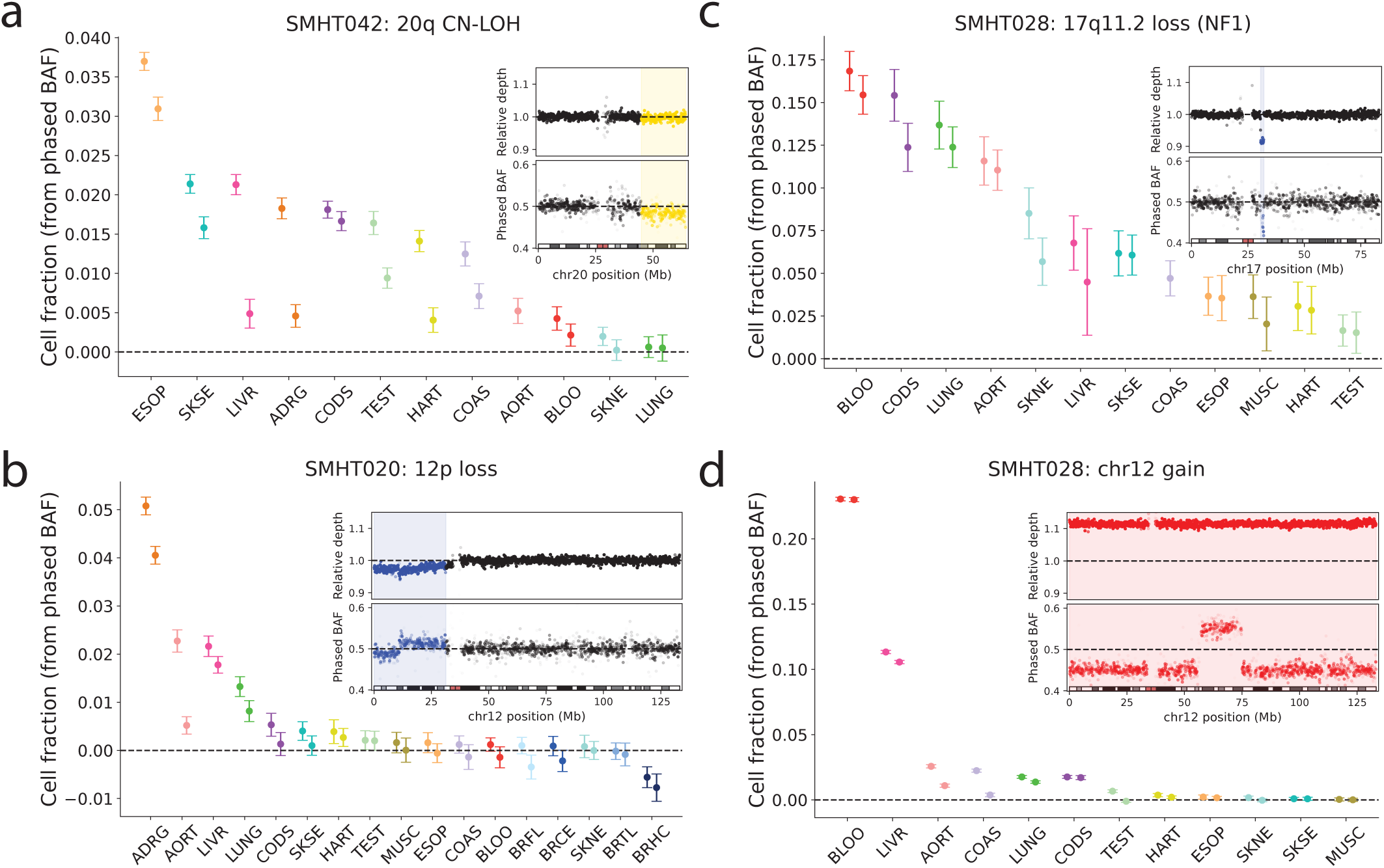
Early developmental mCAs present in adult tissues. (a-d) Estimated cell fractions of mCAs present in many separate tissues from the same donor. In each panel, an index mCA was selected as the mCA call with largest cell fraction among a group of several near-identical mCA calls (same genomic span and same phase) in different tissue cores of a donor. The cell fraction of this mCA in every tissue core from the donor was then estimated based on allelic skew within the mCA region (assuming that the mCA under consideration was the only mCA overlapping the genomic region), accounting for phase switch errors identified within the index mCA call (inset, bottom). Negative cell fraction estimates indicate allelic skew in the opposite direction (presumably generated by a different, overlapping mCA) and were only observed in one tissue of one donor (SMHT020 hippocampus (BRHC), bottom-right of panel **b**). Insets show relative read depth (top) and phased B-allele fraction (BAF, bottom) across the affected chromosome for the tissue core containing each index mCA (esophagus for 20q CN-LOH in SMHT042, adrenal gland for 12p loss in SMHT020, blood for 17q11.2 loss and chromosome 12 gain in SMHT028). Error bars are 95% confidence intervals computed as 1.96 times the standard error of the mean allelic skew across 30 sub-segments of the mCA region.

In addition to these mCAs shared broadly across tissues, several other mCAs were detected in a few tissues, suggesting origins later in development or during aging (**Supplementary Figs. 20 and 21**; **Supplementary Note 2**).

### X chromosome loss shaped by X inactivation and X-linked variation

Another form of early developmental mosaicism—one that occurs in all females—is X chromosome inactivation (XCI), the transcriptional silencing of one X chromosome per cell to achieve dosage compensation^63^. Prior to gastrulation, each epiblast cell randomly inactivates one X chromosome (maternal or paternal) and epigenetically propagates its choice to descendant cells^64^. As a result of this random process, one homolog may more often be inactivated compared to the other in adult tissues^65,66^. XCI skewing has been proposed to subsequently affect skewing of LOX^18^ (because only the inactive X chromosome can be lost without compromising cell fitness^35,38^), but the extent of such an effect has not been evaluated.

The multi-tissue, multimodal SMaHT data set allowed us to estimate homolog-specific levels of X loss and explore factors influencing LOX skewing. Specifically, long-read methylation data enabled measuring XCI skewing and phasing X-chromosome-wide variation (**Supplementary Figs. 22** and **23**), and phased short-read WGS data enabled quantifying, in each tissue, the fraction of cells with loss of each X homolog. Most tissues, in female donors of all ages, exhibited skewed XCI, usually toward the same homolog within a donor^66^ (**Supplementary Fig. 23**). Intriguingly, LOX also typically skewed toward one of each donor’s two homologous X chromosomes (binomial *P*<0.05 for 7 of 9 donors; **Fig. 5a**). XCI skewing was modestly predictive of LOX skewing (*R*^2^=0.18, *P*=1.5 x 10^-8^; Model 1 in **Fig. 5b,c**), suggesting that XCI was one determinant of LOX skewing, but that other influences might also exist.

**Figure 5:**
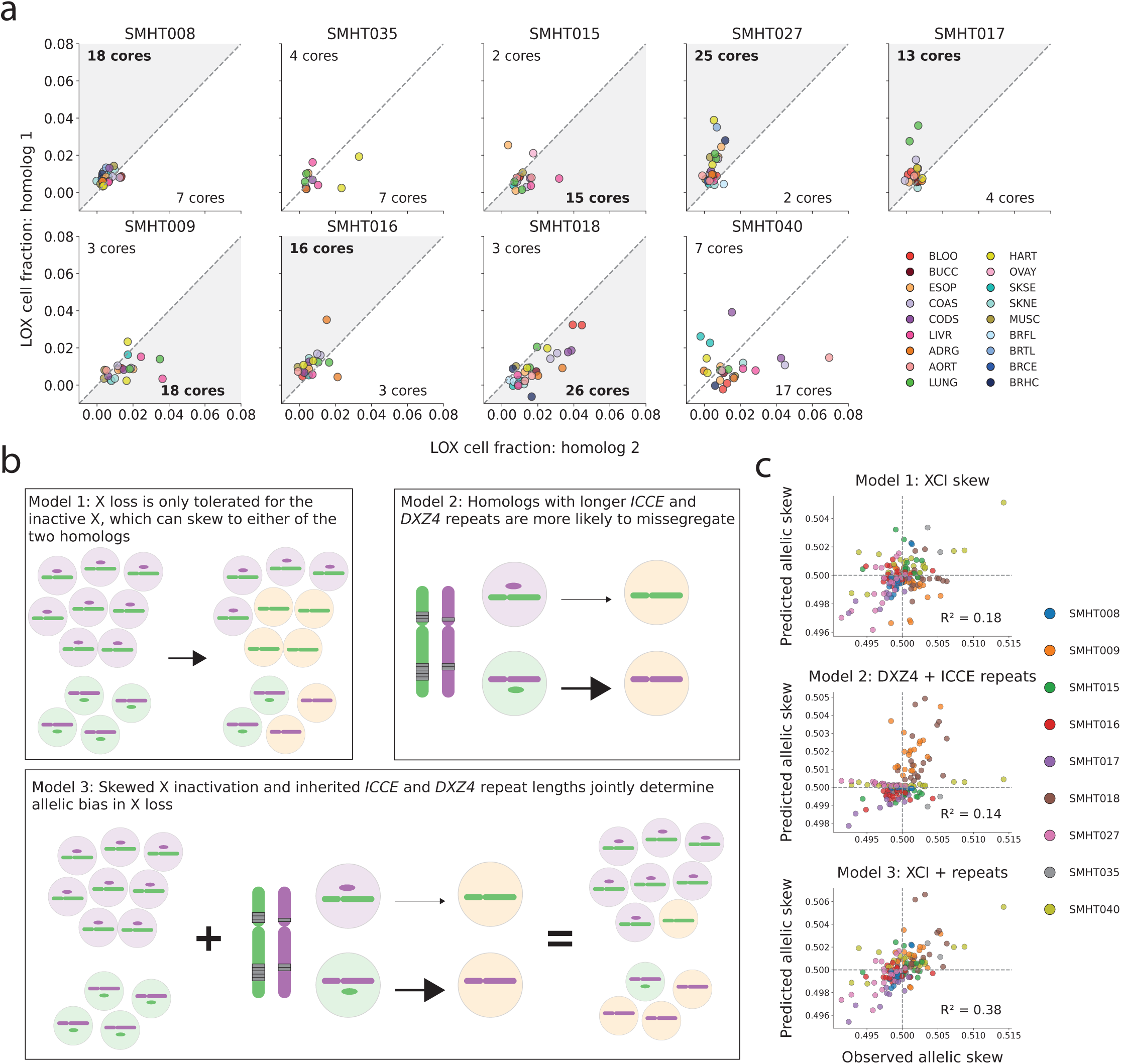
Skewing of female X loss toward one homolog is predicted by X-inactivation skewing and X-linked repeat variation. **(a)** Partitioning of total amounts of X loss in each tissue core into fractions of cells that lost each homolog (1 and 2, corresponding to maternal and paternal in unknown order) based on allelic imbalance at heterozygous sites. Each dot corresponds to a tissue core, with the fraction of cells with homolog 1 lost plotted against the fraction of cells with homolog 2 lost. The number of cores for which homolog 1 (respectively, homolog 2) was lost in a larger fraction of cells is annotated in the top left (respectively, bottom right) of each panel; the label is shown in bold and the corresponding octant is shaded when X loss skews significantly toward one homolog (two-sided binomial *P* < 0.05). **(b)** Models predicting which homolog is more frequently lost in each tissue based on skewing of X chromosome inactivation (Model 1), genetic variation on the X chromosome associated with increased susceptibility to missegregation^35^ (Model 2), or both of these effects (Model 3). **(c)** Scatter plots of the observed allelic skew (allele fraction of homolog 1; x-axes) against the allelic skew predicted by the three models in panel **b**. Each point corresponds to a single tissue core from one of the 9 female donors; data points are colored according to donor. The reported *R*^2^ was computed after randomizing the labeling of homologs 1 and 2 (independently for each tissue core) used when computing allelic skew; this labeling is arbitrary, and randomizing the labeling per core produces a more robust estimate of prediction accuracy in the small-cohort setting.

A distinct factor that could influence LOX skewing is X-linked genetic variation. We recently observed that alleles of two highly polymorphic X chromosomal macrosatellite repeat arrays (*ICCE* and *DXZ4*, which have roles in the physical structure of the inactive X chromosome^67–70^) influence the susceptibility of an X chromosome to loss. Specifically, increased copy numbers of CTCF-binding repeat units in *ICCE* and *DXZ4* appear to reduce an X chromosome’s segregation fidelity^35^. This effect, previously observed in blood, replicated in the pan-tissue SMaHT data set: lengths of *ICCE* and *DXZ4* repeat alleles (which we assembled by combining long-read and short-read data; **Methods**) were also modestly predictive of LOX skewing (*R*^2^=0.14, *P*=1.2 x 10^-6^; Model 2 in **Fig. 5b,c**). Modeling both the XCI and genetic effects together correctly predicted the more-frequently-lost X homolog in 115/161 tissue cores with total LOX exceeding 0.5% cell fraction (*R*^2^=0.38, *P*=3.4 x 10^-18^; Model 3 in **Fig. 5b,c**), demonstrating that female X loss is shaped by independent effects of epigenetic variation in XCI skewing (varying across tissues) and genetic variation between an individual’s two X chromosomes.

## Discussion

Our analysis of mCAs across 19 tissue types of 25 donors demonstrates that healthy adult tissues commonly contain small fractions of cells with aneuploidies and other large mCAs. The patterns in these events revealed characteristics of the mutational and selective forces shaping the landscape of chromosomal mosaicism across the human body. Chromosomal alterations are recurrently generated by mitotic errors but are subsequently filtered by purifying selection, as evident both in age-acquired autosomal aneuploidies—which are typically restricted to a few chromosomes per tissue, often distinct from those frequently aneuploid in cancer—and in the limited (though not uncommon) retention of early developmental mCAs. Notable exceptions are sex chromosome losses, which appear to be fitness-neutral or slightly fitness-increasing across tissues, and aneuploidies in the adrenal gland, which appear to be broadly tolerated. Strong positive selection on mCAs is rare, with the unique exception of the 9q CN-LOH mutations that often arise in the heterochromatic 9q12 hotspot and confer a second hit to *NOTCH1* mutations in the esophagus and skin^21–23^. These factors are broadly distinct from those that shape SNVs and other classes of somatic mutations^34^.

Beyond these chromosome-level and tissue-level propensities for mCAs, we observed evidence of donor-level effects that modulate an individual’s predisposition to sex chromosome loss, including a genetic effect on LOX driven by inherited repeat variation on chromosome X. Based on the strong influence of genetics on mCAs in blood^16–18,35^— explaining ∼50% of variation in hematopoietic LOX and LOY^35^—additional inherited influences on solid-tissue mCAs are sure to be discovered as sample sizes increase. These are likely to include genetic influences on mitotic fidelity and clonal proliferation^35^, including *cis* effects in which mCAs alter copy number of variants influencing clonal fitness^16,71^.

These initial results from 25 donors also point to promising avenues for future work. Increased study sizes and wider sampling of tissues across the full human lifespan will further refine the landscape of mCAs, enabling additional insights into which mCAs are routinely tolerated, when and how cells with mCAs are lost^72,73,61^, and which mCAs experience positive selection too subtle to ascertain here. Increased sample sizes will also eventually allow investigating risk factors (genetic and non-genetic) for solid-tissue mCAs, identifying mCAs associated with clinical consequences^24,8,74^, and evaluating which such associations are causal versus driven by pathology. Emerging single-cell data modalities hold particular promise for identifying and studying mCAs in their cellular and spatial contexts^31,32,57^: whereas our analyses, based on allelic imbalance in bulk WGS data, could not identify cell types containing mCAs or ascertain co-occurrence of multiple mutations within the same cell, future multi-omic studies will be able to observe mCAs together with other somatic mutations within transcriptomically profiled single cells. Collectively, such data will further elucidate a wide-ranging landscape of normal karyotypes and chromosomal variations within healthy tissues.

## Acknowledgements

We are grateful to the SMaHT donors and donor families whose generous gifts have made this work possible. We thank V. Sankaran, S. Raychaudhuri, A. Gusev, and Y. Guo for helpful discussions. This research was supported by the NIH Common Fund, through the Office of Strategic Coordination/Office of the NIH Director under award UM1 DA058230. The All of Us Research Program is supported by the NIH, Office of the Director: Regional Medical Centers: 1 OT2 OD026549; 1 OT2 OD026554; 1 OT2 OD026557; 1 OT2 OD026556; 1 OT2 OD026550; 1 OT2 OD026552; 1 OT2 OD026553; 1 OT2 OD026548; 1 OT2 OD026551; 1 OT2 OD026555; IAA no. AOD 16037; Federally Ǫualified Health Centers: HHSN 263201600085U; Data and Research Center:5 U2C OD023196; Biobank: 1 U24 OD023121; The Participant Center: U24 OD023176; Participant Technology Systems Center: 1 U24 OD023163; Communications and Engagement: 3 OT2 OD023205; 3 OT2 OD023206; and Community Partners: 1 OT2 OD025277; 3 OT2 OD025315; 1 OT2 OD025337; and 1 OT2 OD025276. In addition, the All of Us Research Program would not be possible without the partnership of its participants.

## Author Contributions

D.T. and P.-R.L. conceptualized the project and drafted the manuscript. D.T. performed analyses. N.K. and Y.S. provided guidance on the analyses and interpretation of results. C.E.S., W.C.F., and A.D.V. contributed to data processing at the SMaHT Data Analysis Center under the supervision of H.-J.E.C. and P.J.P. P.-R.L. supervised the project. All authors reviewed and edited the manuscript.

## Methods

### Ethics

This research complies with all relevant ethical regulations. The de-identified sequencing data analyzed here were previously generated by the SMaHT Network, which obtained ethical approval from the National Disease Research Interchange (NDRI) through the University of Pennsylvania (IRB#5 FWA00004028) under NDRI’s Tissue Procurement Program protocol (#704541).

### SMaHT bulk DNA sequencing

Bulk short-read WGS was previously performed at five Genome Characterization Centers (GCCs) on Illumina NovaSeq X Plus and NovaSeq 6000 instruments, and long-read WGS was performed on ONT PromethION (R10.4.1 flow cells) and PacBio Revio instruments. All sequencing data were aligned to GRCh38 (GCA_000001405.15) with ALT contigs and the hs38d1 decoy sequence excluded^75^. Further details of sample collection, preparation, and sequencing are described in the SMaHT flagship paper^34^.

### Tissue nomenclature

For tissues without left/right laterality designations, we used the four-letter tissue abbreviations defined by the SMaHT Network. For tissues that had left/right subcategories, we defined bilateral category abbreviations to indicate merged subcategories: hippocampus (BRHC = BRHL+BRHR), adrenal gland (ADRG=ADGL+ADGR), ovary (OVAY=OVAL+OVAR), and testis (TEST=TESL+TESR).

### Phasing autosomes using statistical phasing and long reads

Subtle allelic imbalances indicative of low-cell-fraction mCAs become much easier to detect when accurate long-range haplotypes are available^15^. We employed a joint strategy leveraging both (i) haplotype sharing with a large reference panel and (ii) physical phasing information from long-read sequencing to produce haplotypes accurate to ∼30 Mb between phase switch errors. In brief, we used a two-step approach that first used statistical phasing to estimate chromosome-wide haplotypes and then corrected apparent switch errors in the statistical phasing using long reads.

Reference-based statistical phasing leverages haplotype frequency information from a large reference panel to probabilistically estimate the relative phase of consecutive heterozygous sites in a target individual^76^. Long reads offer an orthogonal approach to phasing by using physical linkage of alleles on the same read to phase nearby heterozygous sites^76,77^. Unlike statistical phasing, which produces a pair of haplotypes spanning a chromosome that can be probabilistically very accurate when using a large, ancestry-matched reference panel (e.g., ∼10 Mb between switch errors), read-backed phasing produces shorter phase sets (groups of variants with known relative phase) separated by regions that lack phaseable heterozygous variant calls (owing to runs of homozygosity or low mappability). Phase sets are typically 1 Mb or shorter but can be highly accurate if restricted to high-quality variant calls. Thus, phase sets provide an opportunity to correct errors in statistical phasing.

We first performed statistical phasing to generate haplotypes accurate at the scale of ∼10 Mb using the AoU+AnVIL imputation server^78^. The AoU+AnVIL imputation server uses Beagle 5.4^79^ to impute from a reference panel comprising 515,579 individuals of diverse ancestry. We supplied the imputation server with a variant call file containing a merge of the 25 donor-level germline variant calls (generated using DNAscope Hybrid^80^ as previously described^34^, with MLrejected variants excluded). We set the genotypes of variants not called in a donor to hom-REF, and we restricted the imputation input to variants present at >1% global allele frequency in the 1000 Genomes Project^81^. We filtered the resulting imputation output to variants for which the imputed genotype agreed with the called genotype. This set of heterozygous variants (which included successfully imputed rare variants but excluded any variants absent from the imputation panel) formed the statistically phased scaffold on which we performed further phasing refinement using long reads.

Next, for each donor in turn, we performed switch error correction on the phased scaffold by incorporating long-read information as follows:

1. Restrict to SNPs on the statistically phased scaffold passing ǪC filters (i.e., not MLrejected, and with agreement between called and imputed genotypes).
2. Merge all available long-read alignments (across tissues and sequencing platforms), filtering to reads longer than 20 kb with mapping quality greater than 40 and SAM flags 3844 unset. We subsetted to reads longer than 20 kb for computational efficiency as the combined coverage across tissues afforded high coverage even when restricted to the longest, most phase-informative reads (over ∼100x coverage for most donors).
3. Run WhatsHap (v2.8)^77^ with the --distrust-genotypes flag enabled to produce phase sets.
4. For each pair of consecutive variants in the same phase set, set the relative phase of the variants in the statistically phased scaffold to match the relative phase of variants in the read-based phasing.
5. Leave the relative phase of other variants (those not contained in a phase set or those spanning the boundaries of phase sets) unchanged.

This strategy can also be thought of as using the statistically phased scaffold to bridge between read-backed phase sets with the rationale that read-backed phasing gives highly accurate (but fragmented) phase sets.

### Autosomal mCA detection

As in previous analyses, we detected mCAs as extended regions of consistent, phased allelic imbalance in bulk short-read WGS data. We did not incorporate long-read WGS data as the sensitivity of the allelic imbalance-based approach is determined by the total number of sequenced DNA fragments, which is substantially lower for long-read compared to short-read libraries. Our approach built on methods and pipelines that we previously used to analyze blood-derived WGS data in UK Biobank^16^. In brief, our pipeline proceeded as follows: (1) find and mask out germline copy number variants (CNVs) by carefully modeling WGS read depth; (2) count sequenced DNA fragments derived from each of an individual’s two haplotypes, adjusting for technical sources of variant-level bias in allelic balance (REF-bias); (3) call mCAs by modeling allelic imbalance using a hidden Markov model (HMM); (4) assign copy number status to mCAs based on denoised read-depth. While this overall approach followed our previous analysis of UK Biobank^16^, we needed to make several modifications to the pipeline (detailed below) to account for the higher sequencing coverage, presence of multiple samples from the same donor, elevated levels of technical variability in sequencing coverage of solid tissues compared to blood, and limited sample sizes in SMaHT.

First, we computed a single germline CNV mask per donor by analyzing a pooled set of sequencing reads merged across tissue samples. Thus, we masked the same germline CNV regions when calling mCAs from allelic imbalance in each tissue core from a given donor, ensuring uniformity and consistency at the donor level. Second, we adjusted for GCC-specific levels of variant-level REF-bias to account for subtle, but noticeable, differences across sequencing centers. To empirically estimate a variant’s level of REF-bias in each GCC, we aggregated allelic depths across all tissue cores sequenced at the GCC that did not contain an mCA overlapping the variant (using a provisional mCA call set that we generated using REF-bias estimates from UK Biobank^16^); masking out high-cell-fraction mCAs was necessary to prevent signals of genuine, biological allelic imbalance from confounding estimates of technical REF-bias. Third, we modified the HMM parameters to detect lower-cell-fraction mCAs ascertainable from ∼200x sequencing data. Specifically, we discretized the state space to 1/*θ* = 6, 8, 10, 15, 20, 30, 50, 80, 130, 210, 340, 550, 890, 1440, 2330, 3770 where *θ* parameterizes the BAF deviation from 0.5. (As in our previous pipeline^16^, we used discrete values of *θ* in initial HMM analysis and subsequently re-estimated BAF deviations within mCA calls.) Finally, when computing read-depth principal components (which we used to denoise read-depth profiles by eliminating “genomic wave” patterns of technical variation^16^), we pooled per-tissue-core read-depth profiles from female and male donors (across all tissues and all GCCs) and excluded the sex chromosomes in order to maximize sample size. Additionally, we masked provisional mCA calls when computing read-depth principal components to prevent the PCs from capturing genuine, biological variation. We used 20 top PCs from this complete set of per-core read-depth profiles (which captured GCC-level and tissue-level technical variation) to denoise read-depth profiles used to assign copy number status of mCAs (described below).

### Merging over-segmented mCA calls

The HMM-based mCA calling algorithm mostly generated calls that spanned the full lengths of mCAs, but a small proportion of the calls exhibited over-segmentation (i.e., splitting of an mCA into two or more adjacent calls). This mostly affected high-cell-fraction mCAs, within which the HMM sometimes overfit to local fluctuations in allelic imbalance, resulting in the mCA being split into segments, each with a slightly different estimated allelic imbalance. To handle this scenario, we merged consecutive calls separated by 10 or fewer heterozygous sites if their allelic imbalance was within 10% of each other.

After merging, for each mCA call, we computed allelic imbalance (BAF deviation) by summing REF-bias-corrected phased allelic depths across heterozygous sites within the mCA region (accounting for phase switches identified by the HMM). We estimated the standard error of the BAF deviation as the standard error of the mean of the site-level allelic imbalance estimates.

To verify the efficacy of the merging algorithm, we manually reviewed high-cell-fraction mCA calls in the resulting call set (post-merging) and found only one mCA that remained clearly over-segmented. This mCA was a mosaic trisomy 12, called in a liver core from SMHT028, that had been split into four segments. This trisomy 12 probably reflected leukocyte infiltration rather than a “true” liver mCA in hepatocytes (given that SMHT028 had a large trisomy 12 clonal expansion in blood; **Fig. 4d**), so we did not modify the call set to fix this mCA.

### Filtering short mCA calls

In addition to the ∼1,000 mCA calls >1 Mb in length, the initial call set also contained ∼600 calls shorter than 1 Mb. This proportion of short calls was much larger than what we had observed in UK Biobank (in which calls of length <1 Mb constituted only 17% of the call set), and follow-up analyses suggested that the majority of these short calls were false positives. Specifically, stratifying the sequenced tissue cores into higher-noise and lower-noise samples based on variability in 50 kb read-depth profiles showed that the proportion of short calls was a few times larger in higher-noise samples than lower-noise samples. We hypothesize that the spurious short mCA calls reflect technical sources of local allelic imbalance in WGS coverage, which could arise from allele-specific biases in chromatin organization. Such technical variation would not produce artifactual longer calls, as it would neither have consistent magnitude nor consistent directionality (i.e., which haplotype is overrepresented) across longer regions.

In light of the above observations, we conservatively filtered all mCA calls shorter than 1 Mb. We further filtered several short (but >1 Mb) calls at the imprinted 15q11–q13 region, which we previously observed to be susceptible to false positive mCA calls^71^.

Among the 1,008 remaining calls >1 Mb, 64 were 1–2 Mb, 32 were 2–5 Mb, 30 were 5–10 Mb, and the remaining 882 were >10 Mb. The preponderance of large mCA calls and the rapid fall-off in call frequency with increasing mCA length (from ∼600 calls <1 Mb to 64 in the 1–2 Mb range and 32 in the 2–5 Mb range) suggested that the remaining calls >1 Mb were largely true-positive mCAs. The 64 calls in the 1–2 Mb range (most susceptible to technical sources of local allelic imbalance) probably have a higher false positive rate than the remainder of the call set, but these calls clearly also contained real mCAs: one-third of the 1–2 Mb calls were readily attributable to either *NF1* microdeletions across tissues of SMHT028 (12 calls; **Fig. 4c**) or complex karyotype in SMHT040 aorta and colon (10 calls; **Supplementary Fig. 21**).

Beyond these checks on the length distribution of the mCA calls and the characteristics of the shorter calls, we performed several confirmatory analyses that supported the validity of the large majority of the calls (**Supplementary Note 1**). In particular, the several-fold increase in mCA burden from the youngest to oldest donors (**Fig. 1e**) indicated a low baseline rate of false positives comprising at most a small fraction of the calls. Nonetheless, we expect that subsequent analyses of larger cohorts will allow identification of some additional error modes (perhaps at additional interstitial or telomeric locations) and further improve filtering of artifacts.

### Assignment of mCA copy number state

We computed relative read-depth within the boundaries of mCAs called from allelic imbalance to classify events as gains, losses, or CN-LOH. Gains and losses result in increases and decreases, respectively, in read-depth while CN-LOH events have no effect on read-depth. Thus, plotting relative depth versus allelic imbalance separates mCAs into these three classes (**Supplementary Figs. G** and **10**). Unlike our analyses of blood-derived WGS data from UK Biobank, where PC-corrected read-depth was sufficiently accurate to clearly separate even the lowest cell fraction events into distinct classes^16^, PC-corrected read-depth in SMaHT still exhibited substantial levels of technical variation. As such, we adopted a probabilistic classifier to assign copy number state to mCAs while accounting for the high levels of residual technical variation in normalized read-depth.

We started by computing the relative read-depth within an mCA as

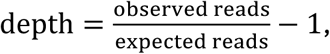

where the expected reads are from the GC-corrected, PC-corrected, aneuploidy-corrected model^16^. We estimated the standard error of this relative read-depth measurement as

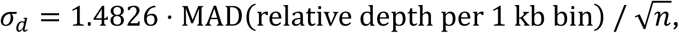

where the MAD (median absolute deviation) was computed across relative read-depth within the *n* 1-kb bins spanned by the mCA.

For each possible mCA class *c* ∈ {GAIN, LOSS, CN LOH}, we then modeled the observed relative read-depth as a Gaussian with mean corresponding to its expectation given the observed allelic imbalance *b* (and assumed copy number class *c*) and variance reflecting both (i) read-depth measurement noise (*σ*^2^_d_ above) and (ii) uncertainty in the allelic imbalance measurement (*σ*^2^_b_), which propagates to uncertainty in the expected read-depth, contributing an additional variance term. Explicitly, we modeled the observed read-depth, conditional on mCA type *c*, as:

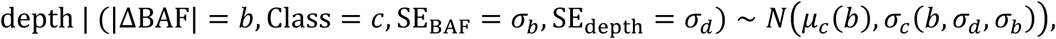

where

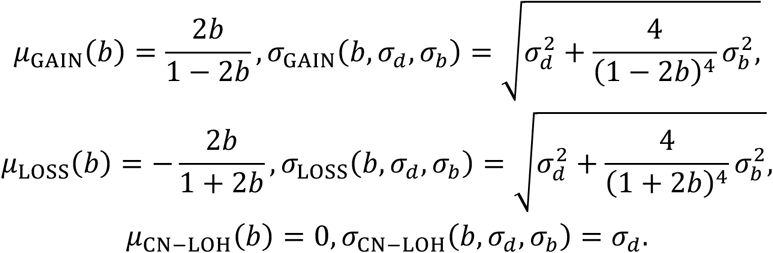

We then used Bayes’ rule to compute posterior probabilities for the three different mCA classes, conditional on the observed BAF deviation and relative depth, and taking into account a prior that captured information about the genomic location and tissue of the mCA call. For our prior, we trained a 3-class logistic classifier to predict mCA type based on chromosome, tissue, and whether the mCA reached the p-arm telomere, the q-arm telomere, or both. We used a subset of 417 mCA calls from samples with low genome-wide read-depth noise (MAD of 50-kb relative read-depth profiles < 0.02) as training data, as these cleaner mCA calls were easier to classify and treat as a ground-truth distribution. In this training data set, calls with relative depth greater than 0.8 times their BAF deviation were treated as gains, calls with relative depth less than –0.6 times their BAF deviation were treated as losses, and calls with relative depth in between were treated as CN-LOH. After fitting the logistic classifier, we then applied it to all mCA calls and regularized the predictions to obtain prior classification probabilities for each mCA: specifically, we computed weighted averages with a uniform distribution across the three classes (80% of the weight on the predicted probabilities and 20% of the weight on the uniform distribution).

We made one further modification to the posterior classification probability computation to prevent overconfidence in classifying mCA calls of intermediate read-depth (between *μ*_LOSS_(*b*) and *μ*_GAIN_(*b*)) that deviated greatly from all three models: for mCA calls of intermediate depth, we clipped z-scores (i.e., (depth − *μ*)/*σ*) in the probability computation to lie between –6 and 6. We only classified calls for which one of the three copy number states had posterior classification probability at least 0.99; as such, intermediate calls for which the z-scores from all three models saturated at |z|=6 were left unclassified. Manual inspection of the classification results showed that this approach produced broadly reasonable results, except for being too conservative in leaving a few mCAs unclassified that had high cell fraction (BAF deviation > 0.02) and read-depth close to but slightly different from zero (|depth| < 0.005). We therefore reclassified these mCAs as CN-LOH, resulting in 902 classified and 106 unclassified mCAs.

### Estimating cell fractions of classified mCAs

We estimated the cell fractions of classified mCAs by combining estimates based on allelic imbalance with estimates based on read-depth. For CN-LOH mutations that do not affect total read-depth, we simply computed their cell fraction as twice the BAF deviation from 0.5. For gains and losses, we averaged their cell fraction implied by allelic imbalance with their cell fraction implied by depth:

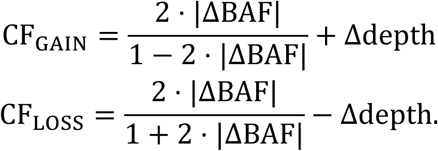

Averaging cell fraction estimates derived from allelic imbalance and read-depth for aneuploidies ameliorated downward bias in allelic imbalance-derived estimates for recurrent mCAs that could potentially affect opposite homologs in the same tissue core, producing competing allelic imbalances.

We did not compute cell fractions for unclassified events as the relationship between allelic imbalance, read-depth, and cell fraction depends on the exact nature of the event.

### Determining relative phase of recurrent autosomal mCAs

Assigning each mCA to a haplotype in a consistent way across overlapping mCAs called in a donor’s tissues was nontrivial due to phase switch errors, which our HMM analyses did not always identify consistently when analyzing allelic imbalance in different tissue cores. Nonetheless, we were able to confidently phase nearly all overlapping mCAs relative to each other based on concordance between allelic imbalance directions within overlapping genomic regions. We represented each mCA as a vector of –1, 1, or 0 corresponding to whether the mCA skewed toward the ALT allele, the REF allele, or exhibited no skew at each heterozygous site on its chromosome (based on the Viterbi path through the HMM). For each mCA, we restricted to its genomic span and computed the cosine similarity between its allelic skew and that of all other overlapping mCAs in the same donor within that span. Since the mCA used to define the genomic span affects the cosine similarity, this produced a pair of asymmetric values for each pair of overlapping mCAs. We restricted to pairs for which the two cosine similarities were either both positive or both negative and recorded the sign of the pair in a symmetric matrix with rows and columns indexing different mCAs.

Next, we assigned each mCA to haplotype 1 (represented by 1) or haplotype 2 (represented by –1) in a way such that the product of the two haplotypes for a pair of mCAs predicted their relative phase (negative if the two mCAs overrepresented opposite haplotypes and positive if they overrepresented the same haplotype). This step was necessary to convert pairwise phase to a consistent global assignment across all mCAs overlapping a given region. Starting from an initial random assignment of mCAs to haplotype 1 or 2, we sequentially chose to flip the haplotype assignments of certain mCAs to minimize the discordance between the observed relative phase and the predicted relative phase. In total, we were able to phase 803 out of 809 phaseable mCAs (i.e., mCAs that overlapped at least one other mCA from the same donor) without error (meaning that all pairwise phases were consistent with a single set of haplotype assignments).

### Identifying mCAs shared across multiple tissues of a donor

In several donors, we observed mCAs with similar breakpoints across tissues, suggesting that some mCAs with shared origins were present in multiple tissues. We classified two mCAs as potentially shared if they exhibited similar breakpoints and affected the same haplotype. More precisely, we collected all pairs of mCAs that affected the same haplotype, resulted in the same copy number, and had genomic spans with a Jaccard similarity >0.9. We then grouped mCAs into sets with potentially shared origins using a union-find data structure where we merged sets containing mCA pairs that were shared. The cohort-wide prevalence of mCA sharing between tissues is depicted in **Supplementary Fig. 20**.

### Estimation of LOX and LOY cell fraction

We recently showed that PC-corrected WGS read depth provides a precise measurement of female LOX and male LOY cell fractions in blood^35^. However, obtaining absolute estimates of LOX and LOY cell fraction using this approach requires accurate absolute calibration of sex chromosomal WGS read-depth: i.e., knowing baseline sex chromosome depths (which can differ slightly from autosomal depths, even after adjusting for ploidy) in individuals without LOX and LOY. In our previous analyses of blood-derived WGS data from biobank cohorts, we obtained this calibration from subsets of several thousand younger individuals (18–22 years old) who had minimal LOX and LOY^35^. In SMaHT, this calibration approach was not possible because only 25 donors were available, with an age range (42– 89+ years) such that nearly all tissue samples from male/female donors harbored appreciable levels of LOY/LOX.

For female LOX, we circumvented this issue by calibrating X chromosomal read-depth in females against X chromosomal depth in males (as LOX in males has been observed to be not tolerated^35,82^). Specifically, we recomputed read-depth PCs in males—now including the X chromosome as well as autosomes—to capture how X chromosomal depth covaries with autosomal depth in individuals unaffected by X loss. That is, we compiled a matrix of read-depth profiles (spanning the autosomes and X chromosome) from all tissue cores of all male donors, and we applied the same read-depth analysis pipeline^16^ to obtain baseline read-depth profiles and top PCs. To denoise X chromosomal read-depth in female tissue cores (and estimate LOX cell fraction), we first adjusted the male-derived baseline read-depth profiles for female X chromosomal ploidy (by doubling the expected read-depth across the X chromosome). We then supplied the ploidy-corrected baselines and the read-depth PCs to the same denoising approach as before^16^, after which we estimated LOX cell fraction based on denoised read-depth across the X chromosome excluding PAR1. We verified that this approach of calibrating against male X chromosomal read-depth agreed with our previous calibration of LOX in AoU blood samples (using 18-to 22-year-old AoU participants)^35^.

To limit the effects of noisy read-depth measurements when comparing LOX levels across tissues and individuals, we excluded tissue cores with high autosomal read-depth variance. For each autosome, we computed the MAD among relative read-depth measurements in 1-kb bins. We then excluded cores with median MAD across autosomes > 0.05. At this threshold, we excluded 40 of 244 total cores (across female donors) but still retained at least one core from each sequenced tissue for all but one donor (for whom both SKNE cores were filtered out). We implemented these exclusions for analyses involving total LOX (**Fig. 2**) but not homolog-specific LOX analyses, which were less sensitive to read-depth noise (**Fig. 5**).

For male LOY, determining baseline Y chromosomal read-depth in samples without LOY was not possible, so instead of estimating LOY cell fraction from read-depth, we used allelic imbalance between the Y chromosome and X chromosome in PAR1 to measure LOY cell fraction. This required knowing the absolute phase of variants in PAR1 (i.e., which allele of each heterozygous site was present on the Y chromosome), which we achieved by combining phase information from long reads and ubiquitous LOY. Compared to the autosomes, PAR1 contains much shorter runs of homozygosity due to its very high recombination rate and greater heterozygosity. As such, long reads from PAR1 tended to produce larger phase sets with few gaps. Among the 16 male donors, long-read phasing on PAR1 produced a median of 2 phase sets (only broken by a large gap in GRCh38 at chrX:1.95–2.13 Mb) and no donor had more than 5 phase sets (**Supplementary Fig. 24**). Due to the high prevalence of Y loss, we were then able to assign absolute phase within each phase set by pooling data across tissues and assigning the haplotype with lower overall allelic depth to the Y chromosome. In each of the donors, we verified that this assignment of absolute phase provided consistent estimates of allelic imbalance across all phase sets on PAR1 (**Supplementary Fig. 24**).

We further verified that LOY cell fractions estimated using allelic imbalance strongly correlated with LOY cell fractions estimated using GC-corrected Y chromosome read-depth measurements (**Supplementary Fig. 4**). We decided to report allelic imbalance-based LOY cell fractions rather than depth-based LOY cell fractions based on the observation that allelic imbalance-based cell fraction estimates exhibited a higher correlation between pairs of tissue cores (Pearson *R* = 0.95) compared to depth-based cell fraction estimates (Pearson *R* = 0.89) (**Supplementary Fig. 3**).

### Testing for donor-level effects on LOX and LOY independent of age and tissue

To quantify the extent to which LOY (and LOX) cell fractions from the same donor were more concordant than expected by chance after adjusting for age and tissue (reflecting donor-level factors such as genetics and exposures), we fit a linear mixed model of cell fraction with fixed effects for age (1 parameter), tissue (1 parameter per tissue), and age × tissue (1 parameter per tissue) and random effects for donor-level effects (1 variance parameter determining the Gaussian distribution from which donor-level effects *d_i_* were drawn) and residual noise (1 variance parameter):

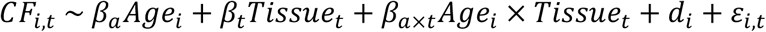

We evaluated statistical significance for nonzero variance of donor-level effects with a likelihood ratio test by comparing with a null model without the random donor effect.

### Calling LOY in cardiac single-nucleus RNA-seq data

In principle, the absence of sequencing reads from the male-specific region of chromosome Y (MSY) provides a way to directly identify nuclei with LOY^39,43,40,83^. However, due to dropout and sparsity, even nuclei that have not lost the Y chromosome will often fail to produce sequencing reads from MSY, especially with low library complexity. As such, we used the following model to assign each nucleus a probability of having LOY, which we then aggregated into cell type specific LOY percentages.

In the model, each nucleus (indexed by *i*) has either retained or lost the Y chromosome, corresponding to latent state *Z_i_* = 0 and *Z_i_* = 1. For each donor *d* and cell type *c*, we place a logit-normal prior on the fraction of nuclei *π_dc_* with LOY:

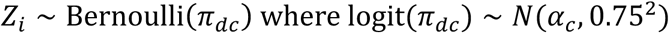

for nuclei *i* from donor *d* and cell type *c*. We further assume that for a nucleus that has retained the Y chromosome, the number of unique molecular identifiers (UMIs) from MSY, denoted *Y_i_*, follows a negative binomial distribution with mean proportional to the total number of UMIs, *N_i_*, and variance parameterized by a cell type specific dispersion factor, *θ_c_*. Nuclei that have lost the Y chromosome are assumed to never produce UMIs from MSY:

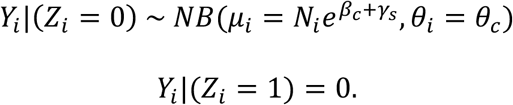

In this parameterization, the variance of the negative binomial component of the mixture is

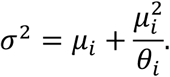

The factor relating the number of total UMIs to the expected number of MSY UMIs for nuclei that have retained chrY, *e^βc^*^+γ^*^s^*, varies according to cell type *c* and study *s* to allow for biological differences in MSY expression across cell types and batch effects due to heterogeneous data sources. The study specific offset, *γ_s_*, is assumed to follow a centered Gaussian prior with variance *σ*^2^ = 0.2^2^.

The posterior probability that a nucleus has lost the Y chromosome is then

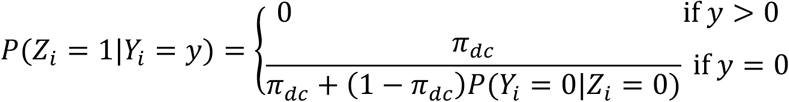

We fit the model parameters by expectation-maximization, using the posteriors to weight nuclei when updating the parameters of the negative binomial model and the prior prevalence. For each donor and each cell type, we estimated the fraction of nuclei with LOY as 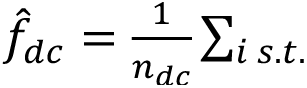 nucleus *i* is of cell−type *c* in donor *d P*(*Zi* = 1|*Yi* = *yi*).

Finally, for each cell type, we estimated the per-decade fold change in the fraction of nuclei with LOY using Poisson regression with robust standard errors to model donor-level probabilistic counts of nuclei with LOY.

### X chromosome phasing using X-inactivation skewing

To facilitate estimating homolog-specific amounts of X loss in female donors, we phased the X chromosome using a custom pipeline integrating statistical phasing, read-backed phasing, and methylation-based phasing. Achieving consistent chromosome-wide phase was uniquely possible for the X chromosome because of the chromosome-wide epigenetic silencing of the inactive X chromosome. When one X homolog is more often inactive than the other, as is often the case in human tissues^66^, the two homologs will exhibit chromosome-wide differences in methylation levels, wherein the more often inactive homolog will have higher bulk methylation levels at CpG islands in promoters that are silenced. Because the SMaHT data set included methylation information from long-read sequencing of DNA from many tissue cores of each female donor, we could combine observed methylation skewing with other sources of phase information to obtain chromosome-wide phase, as detailed below.

For each donor, we generated an initial statistical+long-read-based phasing of the X chromosome using the TOPMed imputation server^84,85^ in conjunction with long reads using the pipeline described above. (The AoU+AnVIL imputation server did not have support for the X chromosome.) In addition to the estimated haplotypes, this pipeline generated a set of high-confidence phase sets for each donor. We next used methylation data to orient these phase sets across the X chromosome, after which we error-checked the results by comparing them to the initial statistical+long-read-based phasing.

For each tissue, within each phase set, we haplotagged reads using WhatsHap and extracted allele-specific methylation at promoter CpG islands (downloaded for GRCh38 from the UCSC Genome Browser^86^) using modkit v0.4.4 (https://github.com/nanoporetech/modkit). A subset of annotated islands did not exhibit methylation patterns consistent with X inactivation (e.g., showing methylation in all reads or no reads), so to remove uninformative CpG islands, we filtered the methylation data to CpGs that had methylation fraction differing by at least 0.01 between the two haplotypes. We then computed allele-specific methylation levels for each phase set as the average number of methylated reads at CpG islands divided by the average coverage (among reads haplotagged with each phase in turn). For each tissue, for each phase set, we computed the signed difference between the methylation levels of the two haplotypes, resulting in a donor-specific matrix with dimensions (number of tissue cores) by (number of phase sets).

By design, long-read-based phase assignments of heterozygous variants were consistent within each phase set but random across phase sets. To determine the relative phase between phase sets, we leveraged the fact that methylation skew within a given tissue should be consistent across phase sets. That is, we wished to identify a subset of phase sets to flip so as to cause the methylation skew within each tissue to agree across phase sets as much as possible. Concretely, we set up an optimization in which each phase set was to be assigned either +1 or –1 depending on whether the haplotype assignments within the phase set agree or disagree with an arbitrary global assignment of haplotype 1 and 2, and each tissue sample was to be assigned either +1 or –1 depending on whether haplotype 1 is more often inactive or active. These two values combine to predict the direction of skew observed in each (tissue, phase set) pair. Specifically, the product of “phase set phase” and “tissue phase” predicts the sign of the methylation skew. We then greedily solved the minimum error correction problem by iteratively flipping “phase set phase” and “tissue phase” to minimize sign disagreements between predicted and observed methylation skews. We treated any (tissue, phase set) pair with absolute methylation skew less than 0.2 (i.e., less skewed than 60% vs. 40%) as missing data to prevent overfitting on tissues with weak X-inactivation skew, and we used 20 different random initializations to avoid local optima. The phase assignments and methylation skew for an example donor are shown in **Supplementary Fig. 22a**.

Finally, we checked the methylation-based assignments of phase set orientation for potential errors by comparing them to the initial statistical+long-read-based phasing. The two approaches showed broadly high concordance (**Supplementary Fig. 22b**). To maximize the accuracy of chromosome-wide estimates of allelic imbalance and X-inactivation skew, we flagged potential phase flip errors (in which 1 or 2 consecutive phase sets locally disagreed within a stretch of phase sets that otherwise agreed between the two phasing approaches) and excluded such phase sets from the computations described next.

### Estimating homolog-specific X loss cell fractions and X-inactivation skewing

Allelic fraction (AF) relates to homolog-specific levels of X loss (cell fractions CF_1_ and CF_2_) as follows:

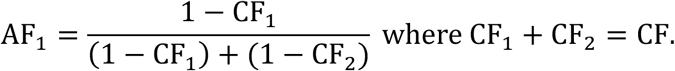

Thus, the homolog-specific X loss cell fractions can be computed from allelic fraction and total cell fraction (CF, computed as 2 ⋅ (1 − relative chrX depth)) as:

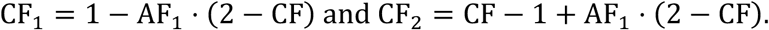

We computed allelic fractions of the two homologous X chromosomes by summing deduplicated allelic depths^16^ (from short-read WGS) across the X chromosome.

We computed the ratio of X-inactivation skew between homolog 1 and homolog 2 as (1 + *M*_1_ − *M*_2_)/(1 + *M*_2_ − *M*_1_) where *M*_1_ and *M*_2_ are the fractions of methylated CpGs (averaged across the X chromosome) on homologs 1 and 2, respectively. The expressions in the ratio come from adjusting *M*_1_ and *M*_2_ to sum to 1 (by adding (1 − *M*_1_ − *M*_2_)/2 to each of *M*_1_ and *M*_2_); in theory, *M*_1_ and *M*_2_ should sum to 1 if CpG methylation perfectly measures X inactivation, but in practice, their sum was typically closer to 0.9.

### Genotyping and phasing *DXZ4* and *ICCE*

*ICCE* and *DXZ4* are large X-chromosomal macrosatellite repeat arrays at which alleles commonly range in length from 0–13 copies of a 5.4-kb repeat unit (*ICCE*) and 10–120 copies of a 2.9-kb repeat unit (*DXZ4*)^35,87^. Given their large size and high intra-allelic homology, these repeat arrays often cannot be spanned by standard long reads and cannot be accurately assembled by standard assembly pipelines. As such, we developed a bespoke pipeline to assemble *ICCE* and *DXZ4* repeat alleles from high-coverage long-read and short-read sequencing data using intra-allelic paralogous sequence variants (PSVs) that distinguished repeat units within and between alleles of female donors.

First, we identified spanning, flanking (i.e., overlapping one flank), and in-repeat long reads containing repeat sequence from *ICCE* and *DXZ4*. To do this, we realigned long reads to a set of reference sequences representing alleles containing homogeneous repeat units of *ICCE* and *DXZ4*. Specifically, the reference contained contigs with between 1 and 20 homogeneous copies of an *ICCE* repeat (chrX: 56770006-56775407 in GRCh38) along with 200 kb of the two flanking sequences surrounding the two near-perfect copies of this sequence in GRCh38, and a contig with 50 homogeneous copies of a *DXZ4* repeat (chrX:115727763-115730717 in GRCh38), along with 200 kb of the sequences flanking the near-perfect copies of this repeat in GRCh38 (most of which are on the minus strand, unlike the reference repeat unit we used, following previous work^35,67^; as such, we reverse-complemented the plus-strand repeat copy extracted from GRCh38). The rationale for including *ICCE* reference contigs of varying lengths was to facilitate alignment and identification of spanning reads (as *ICCE* alleles are sometimes short enough to be spanned by long reads); in contrast, for *DXZ4*, we only included one reference allele (longer than the maximum length of a long read) because *DXZ4* alleles are almost always too long to span^35^. We used *minimap2*^88^ to realign long reads (from all tissues) longer than 10 kb that originally mapped to the *ICCE* and *DXZ4* “bait” regions in GRCh38 (chrX:56767700-56781000 and chrX:115721400-115889220, respectively). We then phased flanking and spanning reads based on SNP genotypes in the flanks. Donors with spanning reads from one or both *ICCE* alleles did not require further assembly of *ICCE* as the combination of spanning reads and diploid copy number (described below) was sufficient to determine the lengths of both alleles. We assembled the remaining *ICCE* alleles along with *DXZ4* alleles using the pipeline described below.

The idea for assembly was to use internal single-nucleotide PSVs, which can be identified from short reads, as landmarks when stitching together in-repeat reads. First, we estimated the diploid copy number for *ICCE* and *DXZ4* based on GC-corrected short-read WGS coverage in blood across the bait regions of GRCh38 as in our previous work^35^. We then used *bwa mem*^89^ to realign all short reads mapping to the two bait regions above to a reference containing 1.5 homogeneous repeat units of *ICCE* and *DXZ4*. We ran a pileup analysis on these realigned reads using *htsbox pileup* (restricting to reads with length ≥50 and positions with base quality ≥20) to count the occurrences of each base at a given position in the reference repeat. We kept variants with VAF between 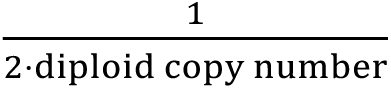 and 0.8 as putative PSVs. For each long-read alignment, we then genotyped each of its repeat units at each of these putative PSVs. We kept only PSVs with both long-and short-read VAF greater than 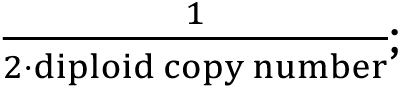 this filtered out spurious PSVs that did not replicate in the long reads. Finally, we represented each long read as a sequence of “barcodes” (one barcode per repeat unit) based on the PSV alleles present within each of its repeats.

To handle occasional long-read sequencing errors at PSV sites, we identified a subset of barcodes that were observed several times in the data set (as would be expected for a genuine repeat-unit variant, given the high long-read coverage merged across tissues) and attempted to assign each repeat unit to one of the barcodes in this set. Specifically, the number of times each barcode was observed in a donor’s long-read data formed a multi-modal distribution with an exponentially decaying mode at 1 corresponding to spurious repeat units containing sequencing errors at one or more PSV sites. For each donor, we filtered out barcodes with counts below a threshold chosen to separate this mode from genuine, repeatedly observed barcodes present in at least one copy in the donor’s genome. We further identified a set of copy number 1 barcodes (forming the first mode of the remaining distribution) representing unique repeat units present in exactly one repeat copy of exactly one of the donor’s two alleles; these served as useful landmarks for assembly.

We next assigned each repeat unit of each long read to its closest-matching barcode that passed ǪC, leaving repeat units unassigned in cases of ties (i.e., matching two or more barcodes equally well). This process of tagging repeat units with closest-matching barcodes provided error-correction, cleaning the long reads used for assembly.

We assembled alleles by starting from the flanks and iteratively extending alleles until extension was no longer possible or until an opposite flank was reached. To allow for the possibility of a phase switch error between the SNPs on the left and right flanks of the repeat, we did not require that the haplotype on the left flank match the haplotype on the right flank (though none of the assemblies ended up containing phase switches). The flanks provided useful seeds for assembly since alignments of flanking reads were unambiguous, and the portions of these reads that overlapped the repeat region provided information about the repeat content (barcodes) of the first several repeat units. For each of the four phased flanks (two on the left and two on the right), our algorithm proceeded iteratively, maintaining the following information at each step:

- A consensus sequence of barcodes representing the current state of the assembled sequence.
- A counts table tracking the number of reads supporting each barcode at each repeat position in the consensus.
- A list of remaining reads available for alignment to the consensus (i.e., not already used).

The overall idea of the assembly algorithm was to progressively incorporate information from long reads with clear alignments to the existing consensus sequence, updating the counts table and consensus sequence with the newly aligned reads at each step, and marking the newly aligned reads as used. The consensus sequence was updated from the counts table after each step by selecting the barcode with the most read support at each repeat position, leaving repeat positions that did not have adequate read support or did not have an unambiguous majority barcode unassigned in the consensus. For each of the four assembly attempts (left hap1, left hap2, right hap1, right hap2), the counts table and consensus sequence were initialized using flanking reads from the seed flank, and the list of available reads was initialized to contain all remaining reads when assembling haplotype 1 and all remaining reads that had not been used in assembling haplotype 1 (from the current direction, left or right) when assembling haplotype 2.

The process of extending the assembly involved iterating between two steps: (1) find the best way to extend the assembly (farther from the seed flank) by identifying reads with the most confident alignments to the frontier of the current consensus, and (2) polish the newly-extended consensus with remaining unused reads that could now be aligned fully within the extended assembly. In the first step, we scored each possible alignment of each unused read that would extend the current consensus by overlap quality (i.e., number of matching barcodes minus number of mismatching barcodes, weighting the latter by barcode similarity). Matches between barcodes determined to occur at copy number 1 were upweighted 5 times relative to matches between other barcodes in the score since they provided unique “landmarks” for alignment within the repeat. The five reads with the highest scores were then used to update the counts table (extending it to include repeat positions spanned by the new reads) and update the consensus. In the second step, all reads with at least one match to the new consensus and at least five times as many matches as mismatches were used to update the counts table and the consensus. This second step was repeated up to ten times to maximally incorporate reads into the assembly before the algorithm returned to searching for new extensions in the first step. This assembly procedure produced four sets of assemblies corresponding to the four different flanks. We verified that the *DXZ4* assemblies from the left and right flank of a given haplotype were highly concordant and that the sum of the lengths of the two assembled alleles matched the diploid copy number estimated from short-read WGS (*R* = 0.95; **Supplementary Fig. 25**).

Finally, we phased the assembled alleles onto homologs 1 and 2 from our X-chromosome-wide methylation-based phasing (described above). This was necessary because the SNP haplotypes we had used for defining flank haplotypes were derived from statistical phasing (to avoid potential drop-out or errors in our primary phasing pipeline at these complex loci). For *ICCE*, we used statistical phasing to determine the relative phase between the SNPs immediately flanking *ICCE* and the nearest phase set in the methylation-based phasing; statistical phasing was highly accurate near *ICCE*, which is located near the centromere where recombination is suppressed and haplotypes are very long^90^. For *DXZ4*, we used a methylation-based phasing approach specific to *DXZ4*. The methylation status of *DXZ4*, like *XIST*, is opposite of the rest of the X chromosome, such that the inactive *DXZ4* is hypomethylated and the active *DXZ4* is hypermethylated^67^. Thus, combined with tissue-level bulk X-inactivation skew, the ratio of hypomethylated to hypermethylated reads from a given haplotype facilitates assignment of each assembled allele to a haplotype. Concretely, in tissues where haplotype 1 is more often inactive compared to haplotype 2, reads used to assemble the allele on haplotype 1 will more often be hypomethylated and reads used to assemble the allele on haplotype 2 will more often be hypermethylated. We classified reads as either hypomethylated or hypermethylated using a two-component Beta mixture model fit to the distribution of methylation levels (defined as fraction of CpGs methylated) across all reads. We verified that the methylation-based phasing for *DXZ4* agreed with a statistical phasing approach (analogous to our approach at *ICCE*) for 8 out of 9 assembled alleles.

### Predicting LOX allelic imbalance from X-inactivation skewing and repeat alleles of *ICCE* and *DXZ4*

We used three models to partition diploid X loss cell fractions (estimated from read-depth across the X chromosome) into predicted homolog-specific levels of X loss. These models used (1) X-inactivation skewing within tissues, (2) repeat alleles at *ICCE* and *DXZ4*, and (3) both sources of information.

For model 1, if *R_XCI_* denotes the ratio of X-inactivation skew between homolog 1 and homolog 2, then assuming that only loss of the inactive X is tolerated, the diploid X loss cell fraction (*CF*) is expected to be partitioned between the two homologs as:

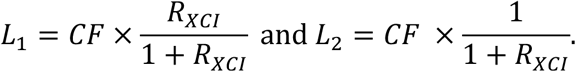

For model 2, for each homolog, we computed the digenic score from Si et al.^35^ to construct a homolog-specific score for the propensity of each homolog to be lost as:

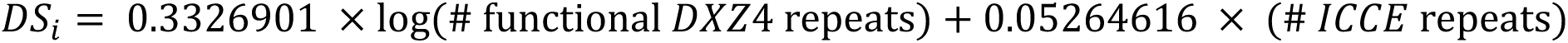

This score is on the scale of log(X loss cell fraction), such that score differential between the two homologs predicts the ratio between the X loss levels of homolog 1 and homolog 2:

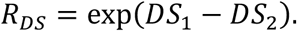

The diploid X loss cell fraction is then expected to be partitioned between the two homologs as:

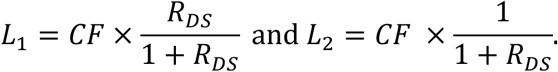

For model 3, the relative propensities for the homologs to be inactive and to missegregate act multiplicatively, such that the homolog-specific levels of X loss partition the diploid X loss cell fraction (*L*_1_ + *L*_2_ = *CF*) according to:

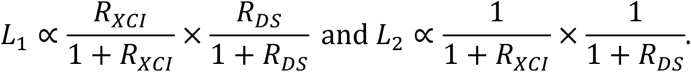

For all three models, the predicted allelic fraction of homolog 1 among retained X chromosomes (reflected in sequencing data) is then:

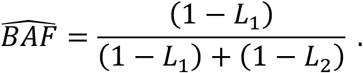

## Data Availability

Sequencing data will be available through the database of Genotypes and Phenotypes (dbGaP) (https://dbgap.ncbi.nlm.nih.gov/) under the study accession number phs004194. Autosomal mCA calls and sex chromosome loss cell fractions are deemed open-access data as per SMaHT Data Use Policy (https://smaht.org/data-use-policy/) and will be available on the SMaHT Data Portal (https://data.smaht.org) prior to publication. More information is available at the SMaHT Data Portal at https://data.smaht.org/.

## Code Availability

The pipeline for calling autosomal mCAs in WGS data is available at https://github.com/tangdavid/mCAs_WGS. Custom analysis scripts will be deposited in a Zenodo repository upon completion of peer review.

