## Supplementary Material for "Recurrent mitotic error and purifying selection shape chromosomal mosaicism across human tissues"

### Supplementary Note

#### 1. Evidence supporting robustness of mCA calls

We identified multiple lines of evidence supporting the robustness of autosomal mCA calls and LOX and LOY cell fraction estimates. These lines of evidence included consistency of results with expected biological patterns and internal consistency of features of the short-read WGS data.

**Biological evidence:**

- **Age effects.** The burden of autosomal mCAs (mean calls per tissue core) increased markedly with age (Fig. 1e), to the extent that this age effect could even be observed for individual tissues with high mCA burden (Supplementary Fig. 6). Similarly, median amounts of X loss in females and Y loss in males increased with age (Supplementary Fig. 8). These patterns were consistent with age-related mCA accrual, as opposed to technical artifacts, which would be uncorrelated with age. Importantly, the several-fold difference in autosomal mCA burden between the oldest and youngest donors provided strong evidence of effective false-positive control (by bounding the baseline rate of false positives at a level well below the cohort-wide call rate; Fig. 1e). This pattern was also clear for male LOY burden (Supplementary Fig. 8).
- **Tissue specificity.** The genomic distributions of autosomal mCAs were very different for different tissues (Fig. 3a and Supplementary Fig. 15). This tissue specificity is consistent with differing selection for or against specific mCAs in different tissues and would not be expected from false positives (e.g., germline CNVs or technical artifacts, which would tend to be broadly observed across many tissues).
- **Consistency with prior literature.** The autosomal mCAs most commonly called at high cell fractions in SMaHT (9q CN-LOH in skin and esophagus and aneuploidy in adrenal gland; Fig. 1f) were also the most prominent events in the prior pan-tissue analysis of GTEx.[1] Additionally, the single-cell analysis of 13 adrenal gland samples in Gao et al.[1] found one sample with a highly expanded clonal population of cells with multiple aneuploidies, consistent with the general pattern observed in SMaHT (Fig. 3c).

**Technical evidence:**

- **Consistency of read-depth and allelic imbalance.** Read-depth deviations within autosomal mCA calls (made using only allelic imbalance between an individual's two haplotypes) were broadly consistent with the calls reflecting the three categories of mCAs (gain, CN-LOH, and loss), with relative abundances varying by tissue (Supplementary Fig. 10). Among whole-chromosome autosomal aneuploidy calls, read-depth deviations in p-arms closely matched those in q-arms (Supplementary Fig. 11). For LOY, allelic imbalance measurements in the PAR1 pseudoautosomal region were consistent with read-depth deviations in male-specific Y chromosomal regions (Supplementary Fig. 4).

- **Chromosome-wide phase consistency.** For donors for which multiple overlapping autosomal mCAs were called on the same chromosome (in different tissue cores), these calls largely agreed on the same chromosome-wide phasing of the donor's two homologs (i.e., phase switch errors detected by the HMM were found in the same places across different cores). This indicated that the allelic imbalances from which mCA calls were derived respected haplotype structure, as expected for genuine mCAs but not technical artifacts.

- **Consistency across sequencing centers.** Samples sequenced at different centers showed broadly consistent tissue-specific burdens of autosomal mCAs (Supplementary Fig. 26).

We also considered directly validating mCA calls using orthogonal technologies (e.g., long read sequencing or RNA-seq) but concluded that this would not be informative. The main problem is that unlike blood, which is a well-mixed system in which different sample aliquots assayed using different techniques provide independent information about the same mCAs,[2] solid tissues are spatially heterogeneous, such that different specimens often contain cell populations with distinct somatic mutations (as evidenced by limited overlap between SNVs found in different cores[3]). As such, a low-cell-fraction mCA detected from short-read WGS of one tissue core would not be expected to be found in a different tissue core (Supplementary Fig. 5), which would be needed for direct validation using another technique, as the original tissue core was already consumed by sequencing. Orthogonal validation could in theory be attempted for the small minority of high-cell-fraction mCAs, but given that these mCAs already strongly matched patterns observed in previous studies, we did not pursue further validation beyond the robustness analyses described above.

#### 2. Example of a clone with complex karyotype found in three different tissues from the same donor

Most mCAs were only detected in a single tissue from a given donor, consistent with late-arising, lineage-constrained events (Supplementary Fig. 20). However, in addition to the early developmental events and the blood-derived trisomy 12 described in the main text (Fig. 4 and Supplementary Fig. 19), we found one other case of a donor (SMHT040) with several mCAs shared across three cores from the abdominal aorta, descending colon, and ascending colon (Supplementary Fig. 7). These three cores contributed 53%, 70%, and 56% of all calls in the aorta, descending colon, and ascending colon, respectively. Moreover, over half of the mCAs called in each of these three cores were found in at least one of the other two cores.

Upon further inspection, we noticed that the allelic imbalance and read-depth profiles from these three tissue cores were highly abnormal, often containing several segments with varying levels of allelic imbalance within a single chromosome (Supplementary Fig. 21). Furthermore, unlike simple mCAs that follow a well-defined relationship between allelic imbalance and depth, mCAs detected in these three cores often exhibited allelic imbalance and depth that did not cluster neatly into gains, losses, and CN-LOH mutations (as evidenced by the clusters of unclassified mCAs in AORT, CODS, and COAS in Supplementary Fig. 10). This evidence points to the presence of a clone harboring a highly abnormal karyotype, akin to a cancer genome. The origin of this clone is not entirely clear as these anatomical sites are not proximal. However, clonal SNVs detected in the same three cores concurred with the mCA data, suggesting that such a clone does exist and the calls do not reflect a technical artifact.

### Supplementary Figures

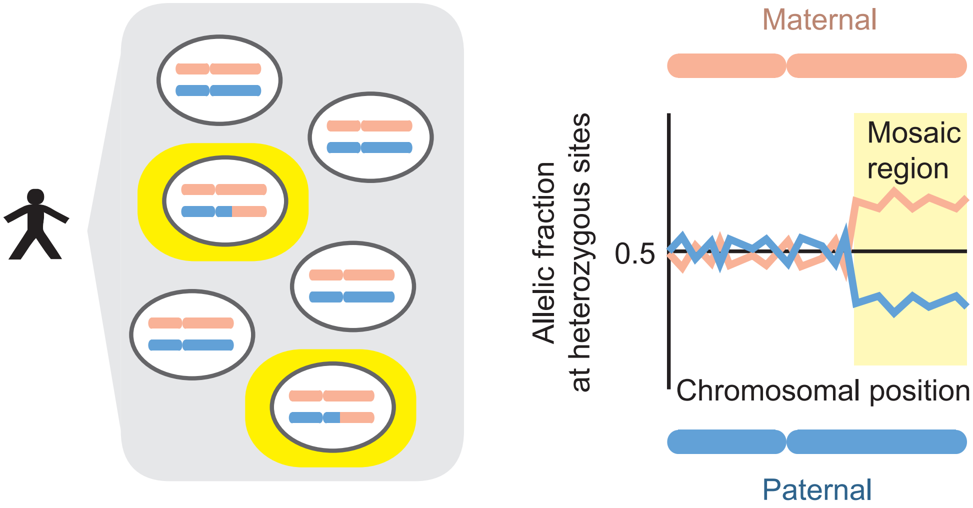


**Supplementary Figure 1: mCAs generate consistent allelic imbalance in bulk sequencing data**. Populations of cells with chromosomal alterations (including gains, losses, and CN-LOH) cause one haplotype to consistently sequence deeper than the other over a large genomic region.

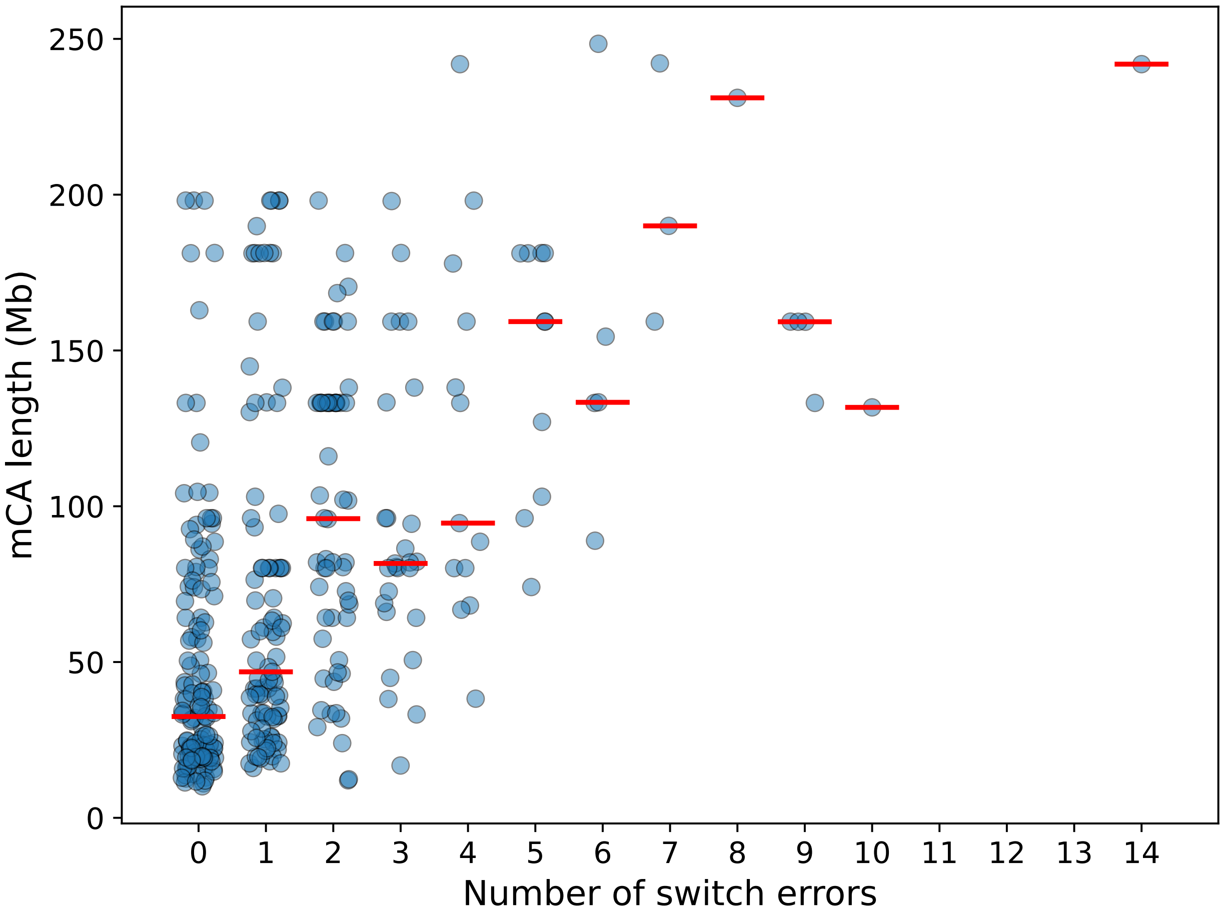


**Supplementary Figure 2: Long range phase accuracy as measured by switch errors in mCAs**. Event length is plotted against number of switch errors detected within each mCA call larger than 10 Mb with a cell fraction greater than 1% (such that most switch errors are detectable by HMM analysis of allelic imbalance). The red horizontal lines indicate median mCA lengths among mCAs with a given number of switch errors.

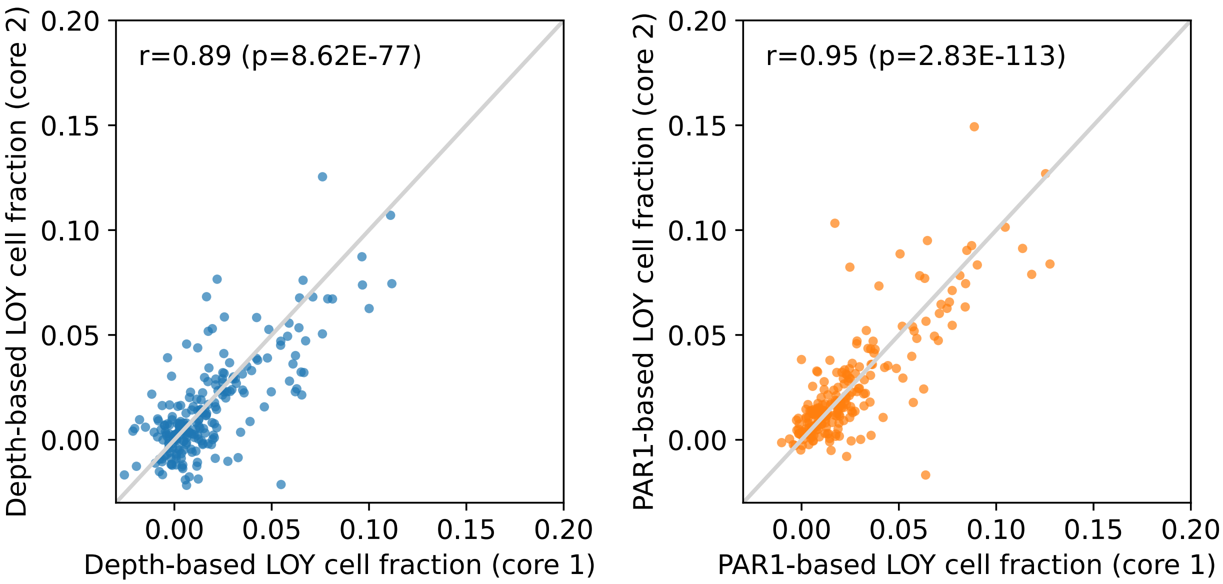


**Supplementary Figure 3: LOY cell fraction in solid tissues is more accurately estimated by allelic imbalance in PAR1 than read-depth in non-pseudoautosomal regions**. Cell fraction estimates from two different cores from the same tissue in the same donor are plotted against each other for read-depth-based estimates of Y loss cell fraction (left) and PAR1 allelic imbalance-based estimates (right). Each point corresponds to a pair of cores (chosen randomly in the rare instances in which three cores of a tissue were sequenced). A few outliers from clonal expansions of LOY in the blood are outside the plot range and not shown.

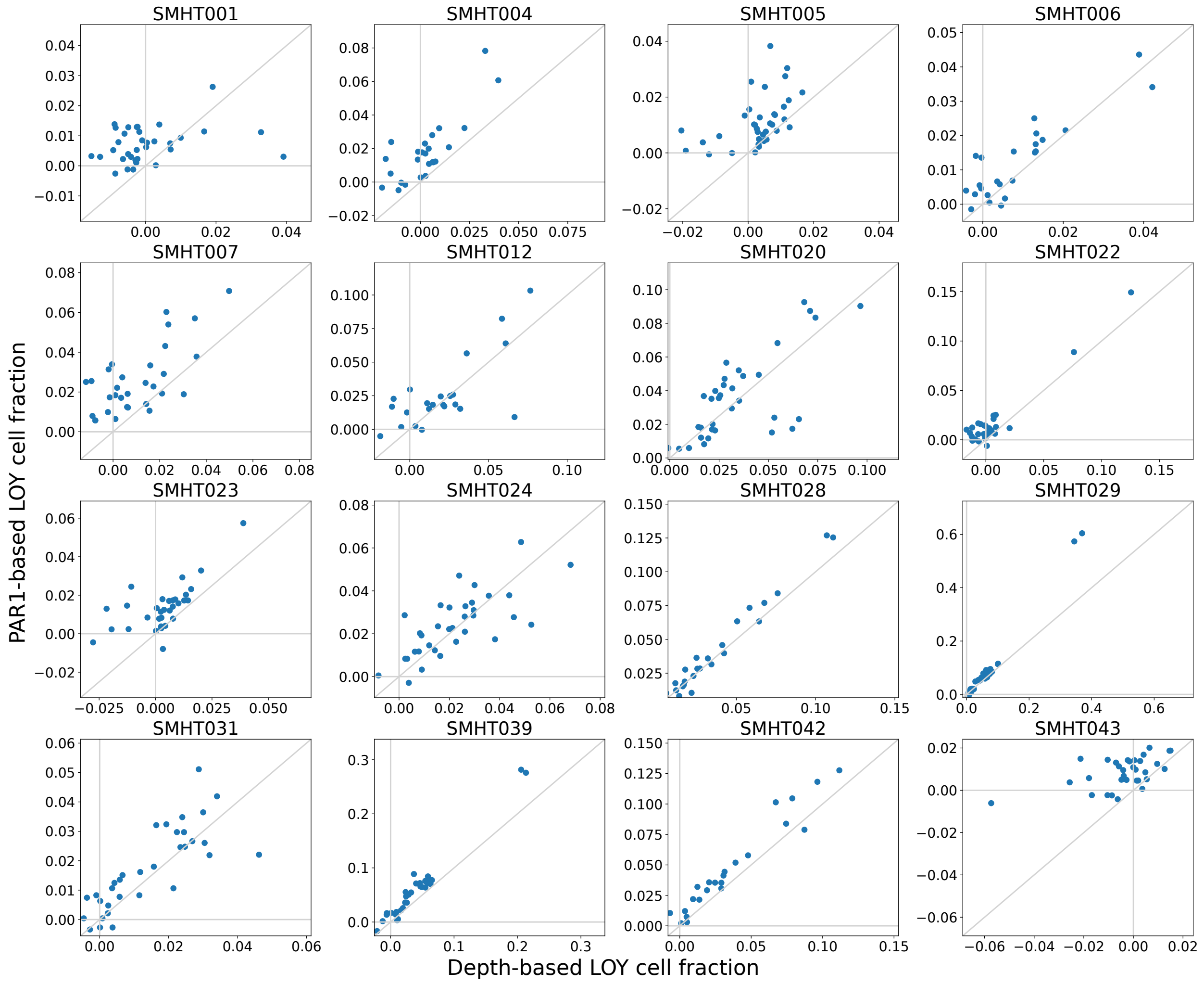


**Supplementary Figure 4: Estimates of LOY cell fraction from Y chromosomal read-depth and allelic imbalance within PAR1 are broadly consistent.** Allelic imbalance-based estimates of LOY cell fraction (y-axis) are plotted against read-depth-based estimates (x-axis) for all tissue cores from all male donors. The two orthogonal measurements offer independent evidence of high levels of LOY observed in solid tissues.

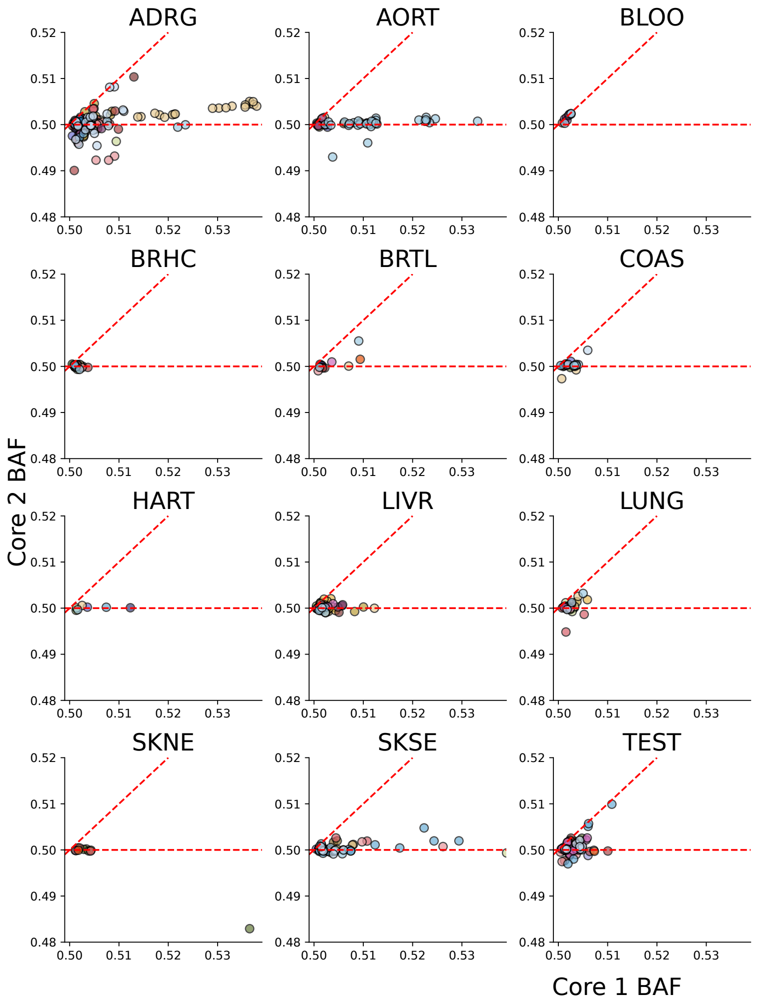


**Supplementary Figure 5: Most autosomal mCAs are specific to a single core.** Allelic imbalances for mCAs ascertained in one tissue core (core 1) are plotted against allelic imbalances over the same genomic spans in a second tissue core (core 2) from the same donor. Switch errors are corrected for according to the ascertained mCA (see Fig. 4). The limited correlation between the two measures of allelic imbalance suggests that most autosomal mCAs are only found in a single tissue core. Only autosomal mCAs greater than 20 Mb are shown. High cell fraction events outside the plot range are not shown. Points are colored by donor.

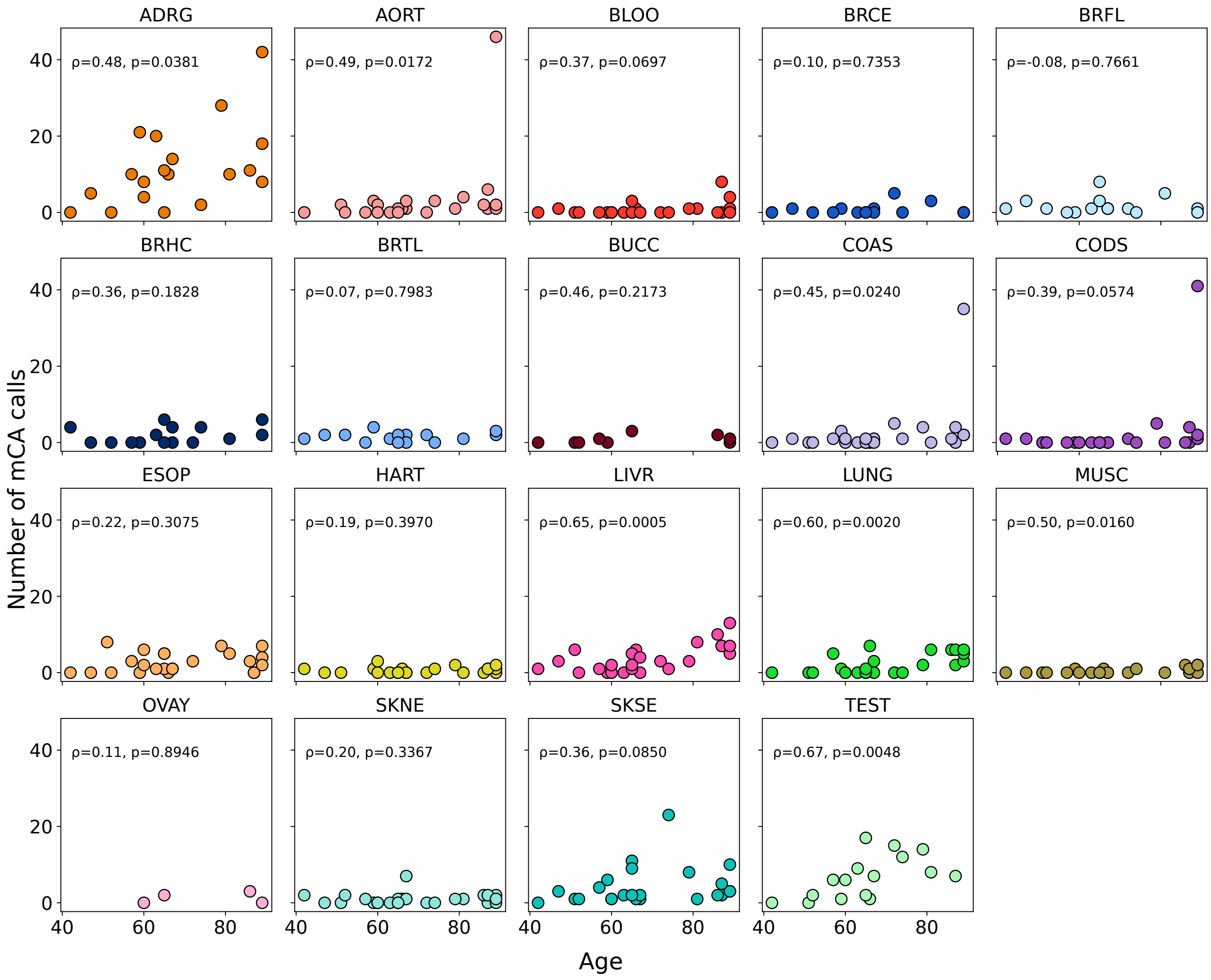


**Supplementary Figure 6: Relationship between autosomal mCA burden and age, stratified by tissue**. Each panel shows the relationship between mCA burden and age in a given tissue. For several of the tissues with notable mCA patterns highlighted in the main text (ADRG, TEST, LIVR, and LUNG), a significant association was observed between mCA burden and age, supporting the validity of the mCA calls.

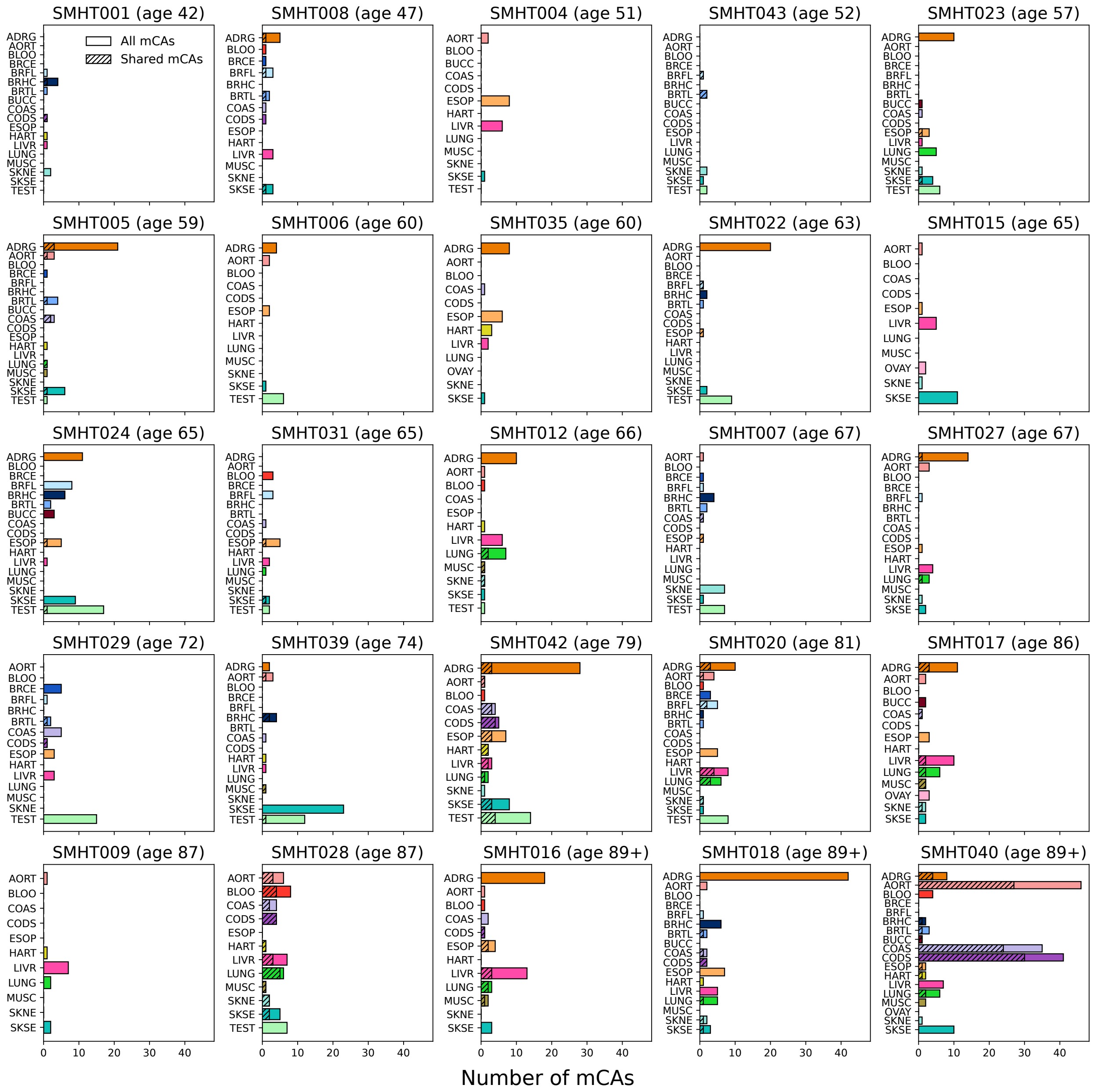


**Supplementary Figure 7: Per-tissue counts of autosomal mCAs called in each donor**. The shaded region of each bar indicates the fraction of mCAs from that particular tissue that are shared across multiple tissues. Donors are sorted left to right and top to bottom by age.

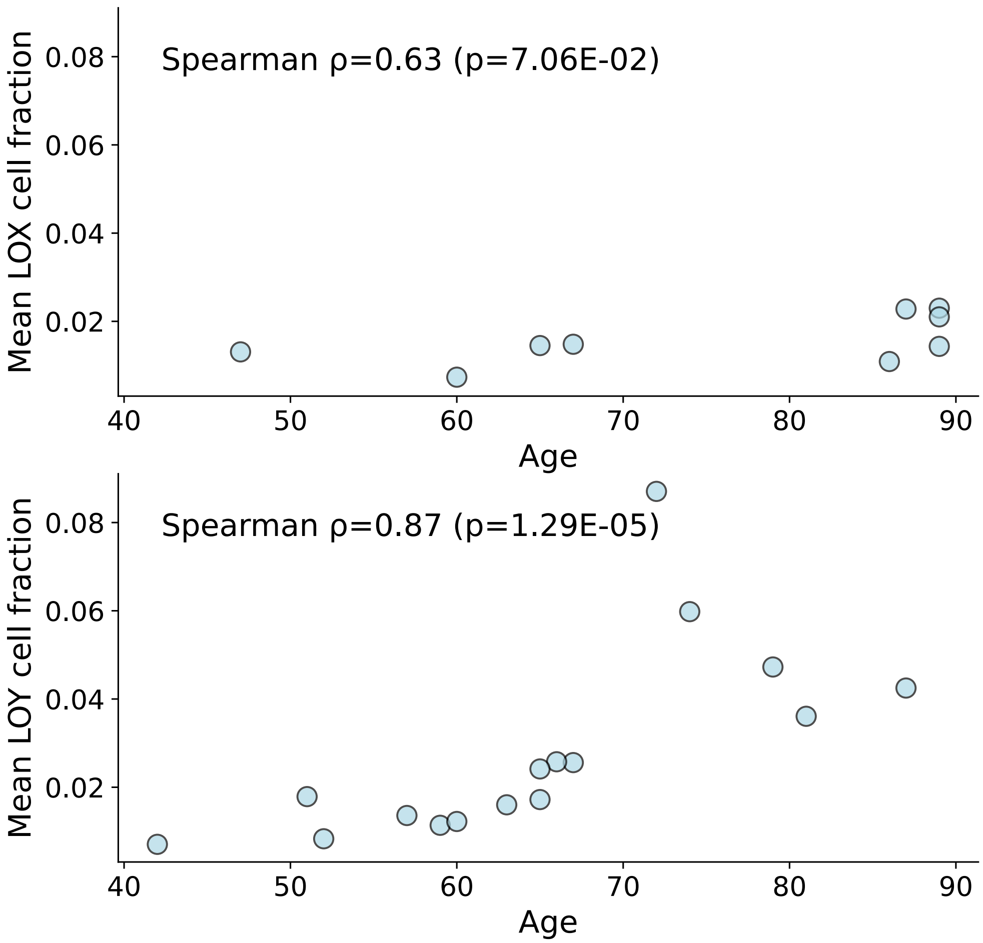


**Supplementary Figure 8: Female LOX and male LOY increase with age**. The mean LOX (top) and LOY (bottom) cell fractions across all tissues from a given donor are plotted against the donor's age.

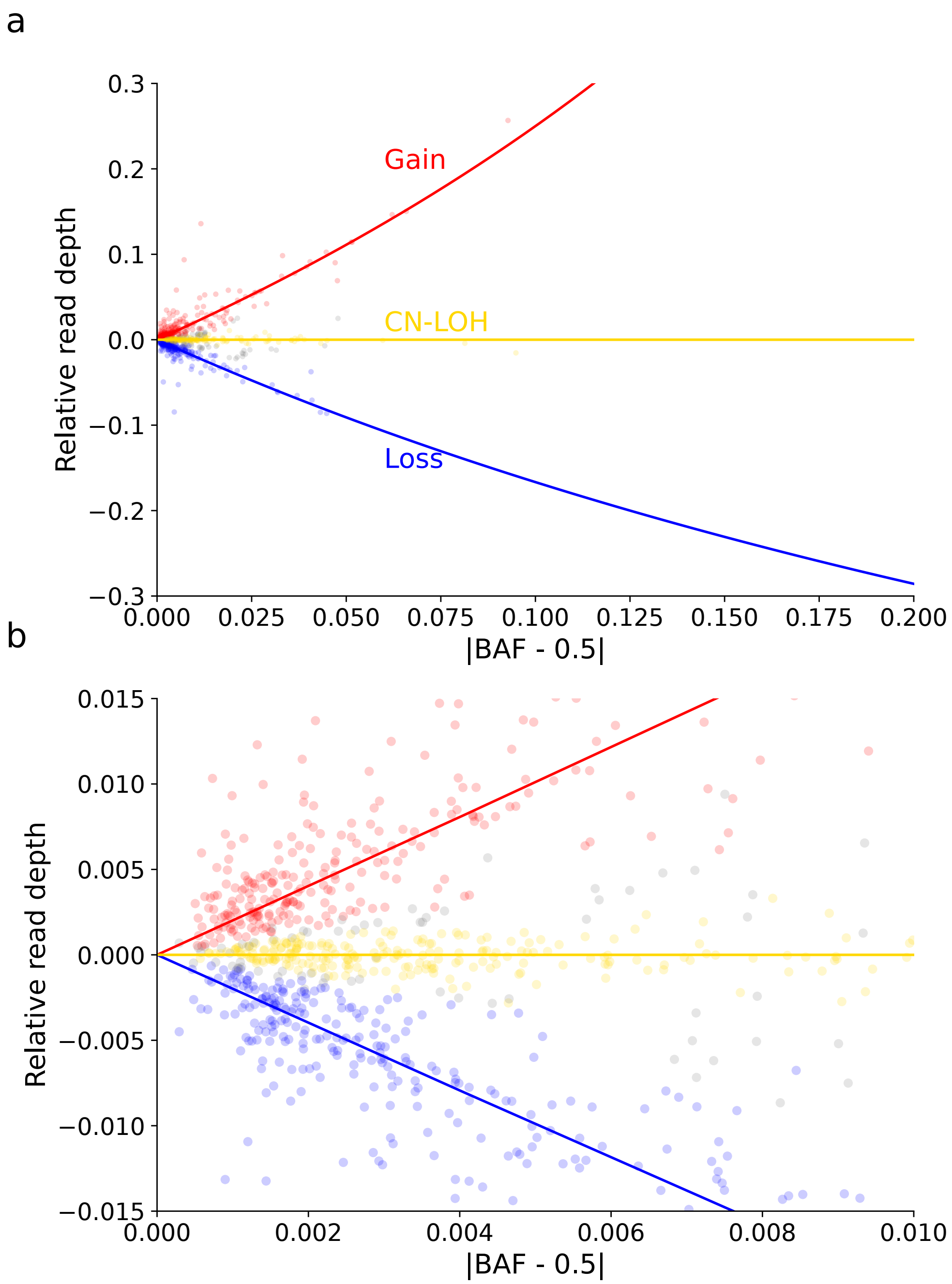


**Supplementary Figure 9: Assignment of autosomal mCA copy-number state based primarily on allelic imbalance and read-depth**. Allelic imbalance and read-depth deviation (i.e., fractional increase or decrease relative to expectation assuming diploid copy number) for called mCAs cluster along three curves corresponding to gains (increased read-depth), CN-LOH mutations (neutral read-depth), and losses (decreased read-depth). Each mCA was assigned a copy-number state using a probabilistic model incorporating information about genomic location and tissue (Methods). Panel **b** is a zoomed-in plot that demonstrates separability of clusters at low cell fractions.

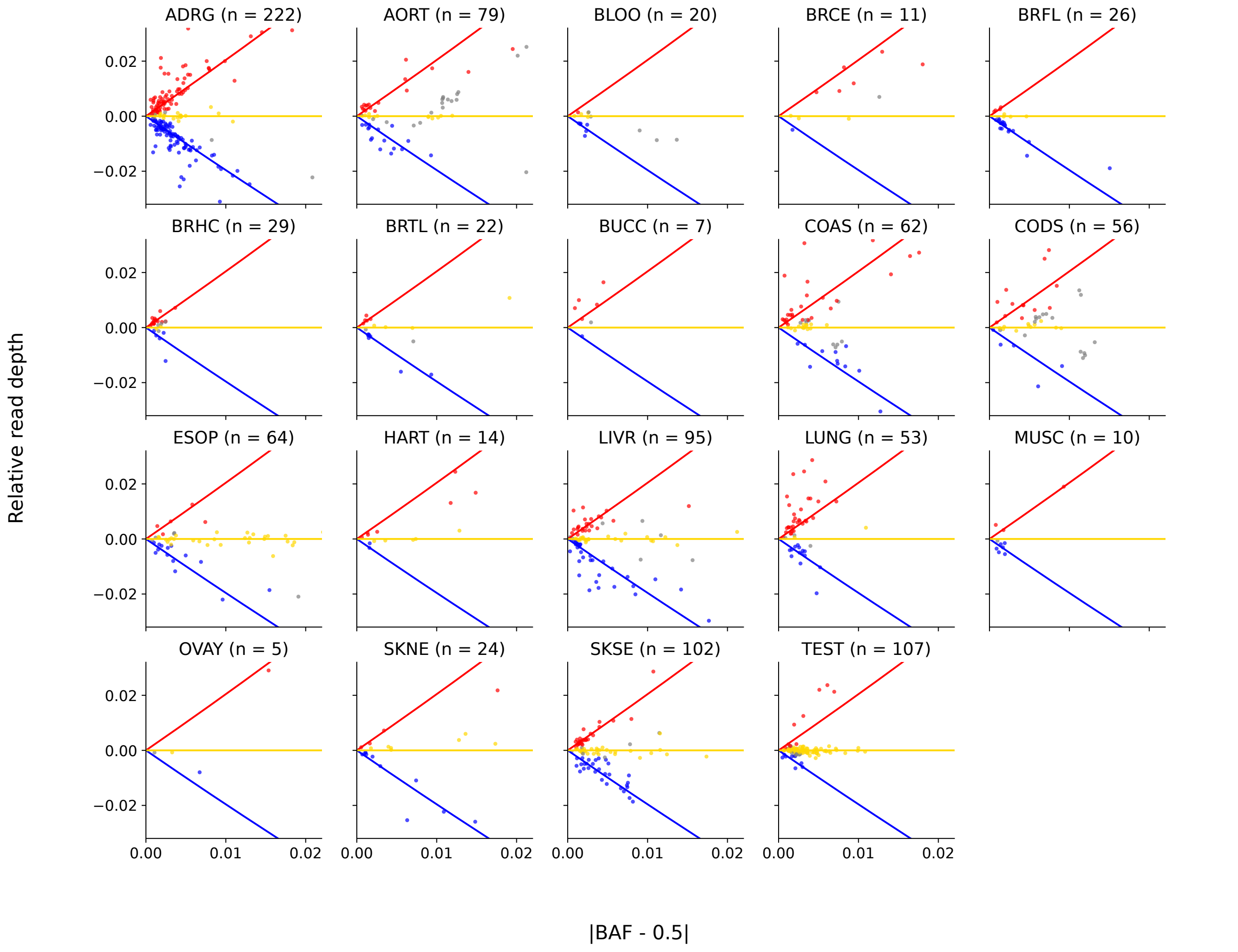


**Supplementary Figure 10: Allelic imbalance and read-depth of mCAs, stratified by tissue**. Each panel shows the relationship between allelic imbalance and read-depth for autosomal mCAs called in a given tissue. Some high-cell-fraction mCAs (such as CN-LOH in esophagus) are outside the plotted range and are not shown.

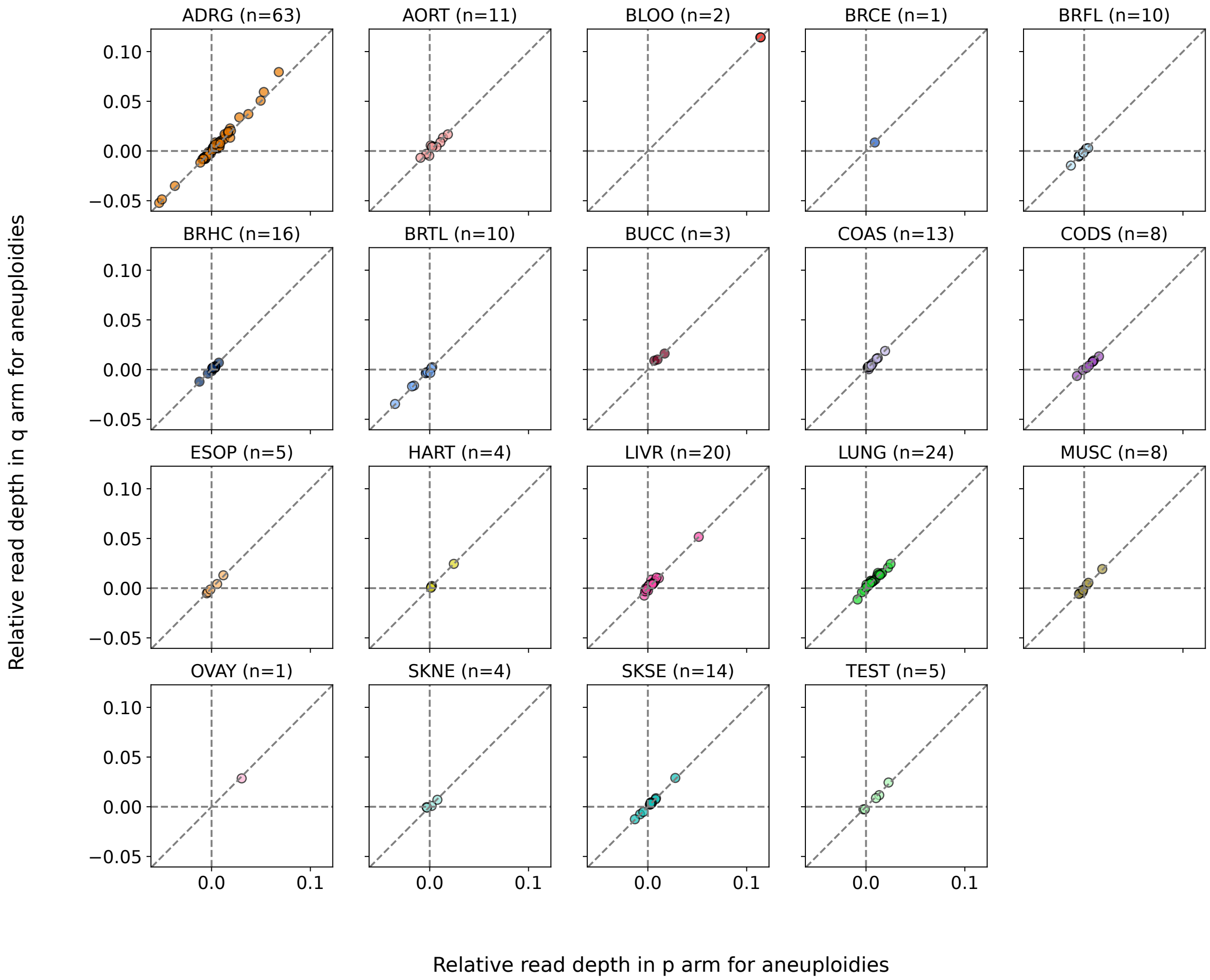


**Supplementary Figure 11: Consistent read-depth in the p and q arms of whole-chromosome aneuploidies**. Read-depth measurements in the p-arm are plotted against read-depth measurements in the q-arm for autosomal aneuploidies called across regions spanning at least 90% of a chromosome. Events on acrocentric chromosomes were excluded.

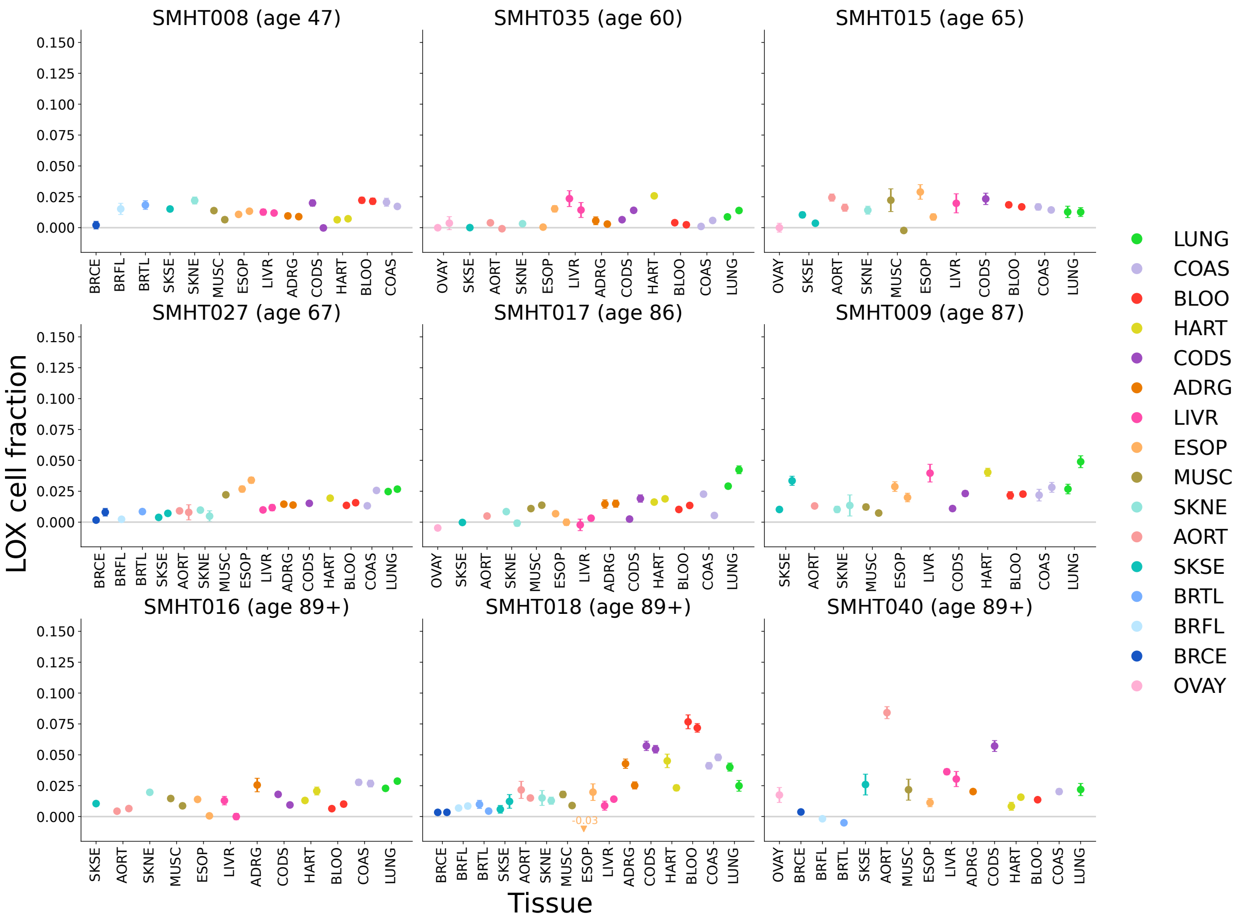


**Supplementary Figure 12: Pervasive age-associated X loss across tissues of female donors**. Estimated fractions of cells with loss of X in each available tissue core from all female donors, sorted by age. Tissues are sorted by expected LOX fraction at age 70 based on linear regression of cell fraction on age. Error bars are 95% confidence intervals.

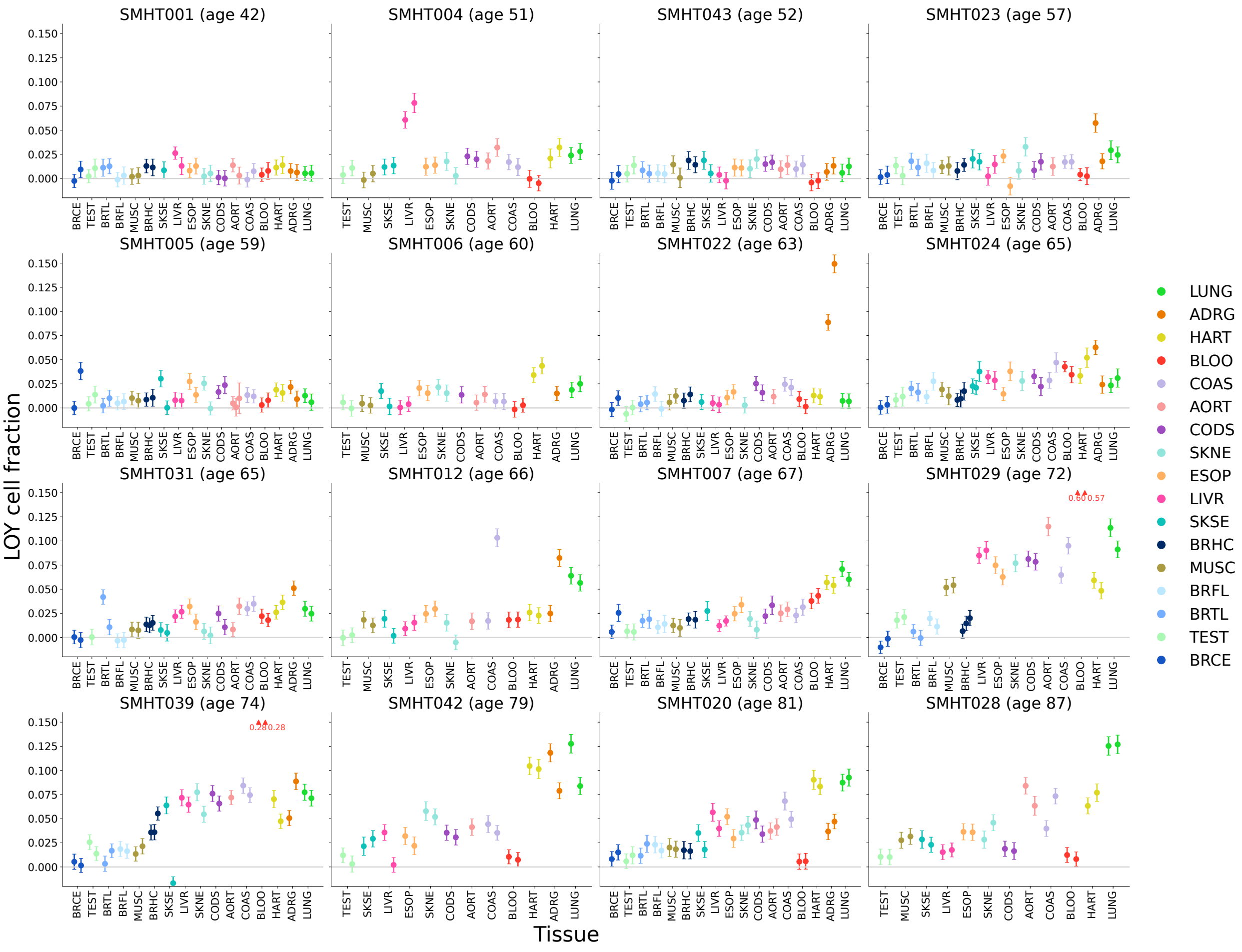


**Supplementary Figure 13: Pervasive age-associated Y loss across tissues of male donors**. Estimated fractions of cells with loss of Y in each available tissue core from all male donors, sorted by age. Tissues are sorted by expected LOY fraction at age 70 based on linear regression of cell fraction on age. Error bars are 95% confidence intervals.

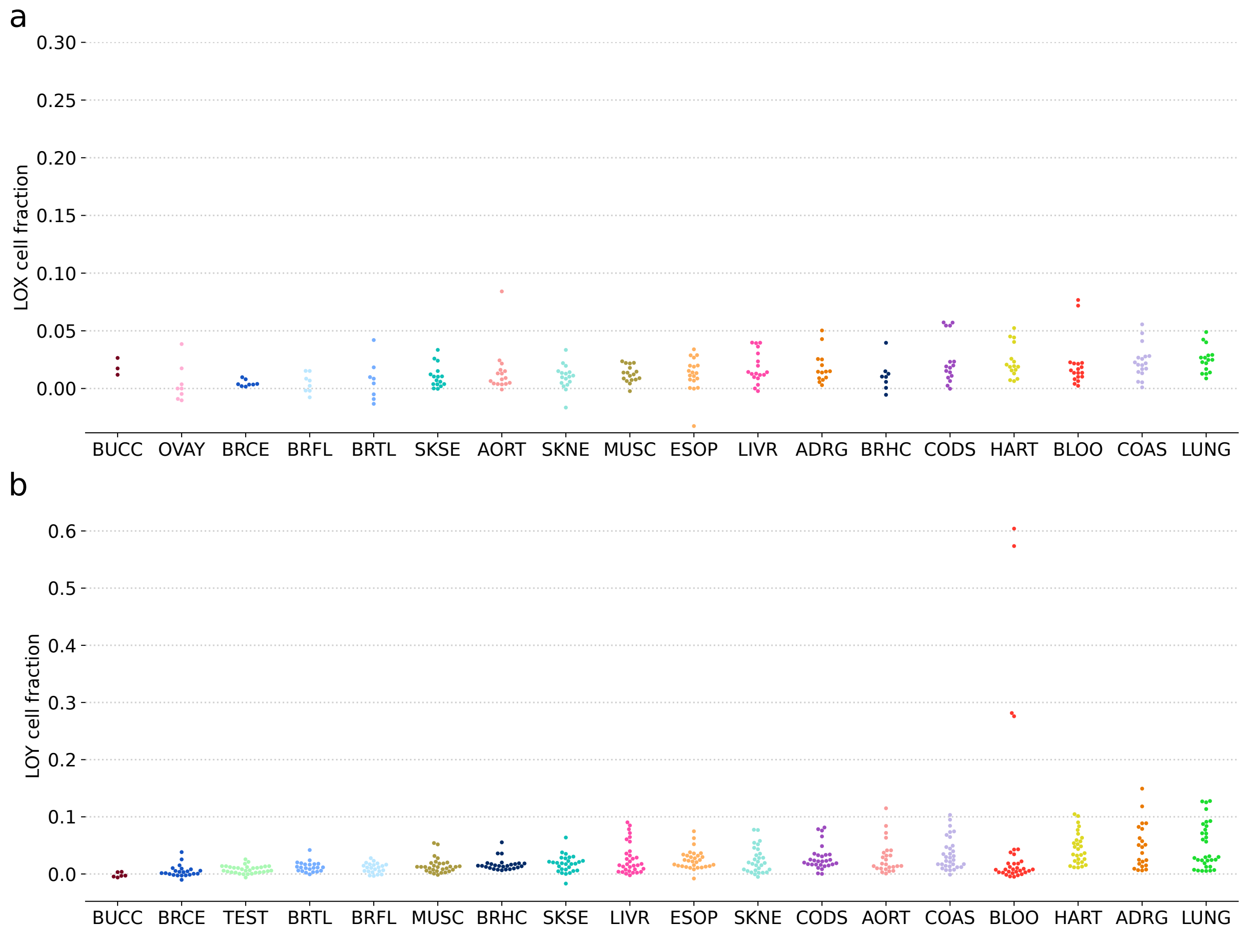


**Supplementary Figure 14: Tissue-level summaries of female LOX and male LOY cell fractions**. **(a)** LOX and **(b)** LOY cell fractions in tissue cores across the SMaHT cohort, grouped by tissue. Tissues are sorted by expected LOX (respectively, LOY) cell fraction at age 70 from linear regression of cell fraction on age.

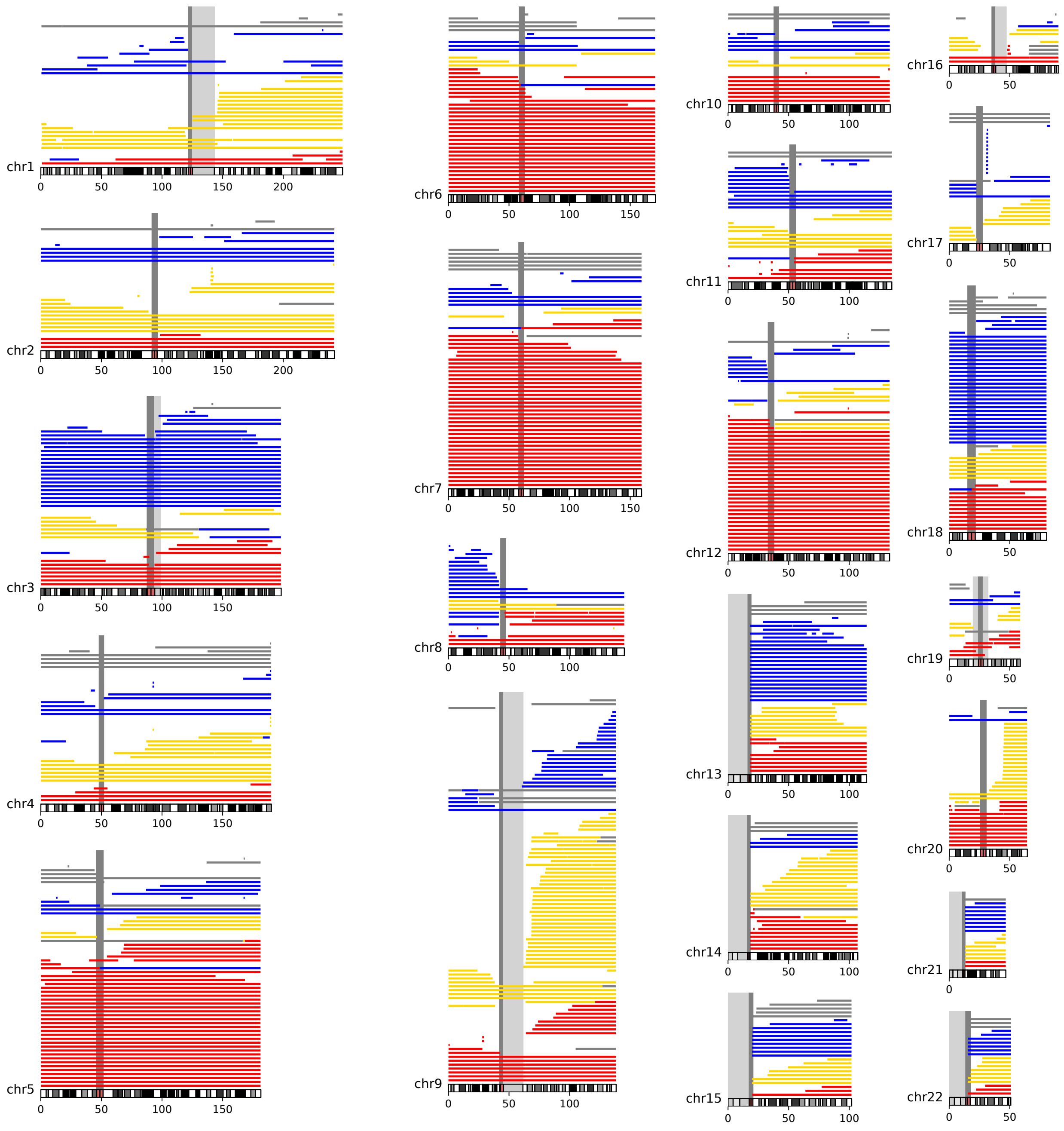


**Supplementary Figure 15: Genomic distribution of autosomal mCAs**. Each line depicts the genomic span of one mCA with mCAs from the same tissue core plotted at the same y-coordinate. Lines are colored by copy-number state: loss, blue; CN-LOH, yellow; gain, red.

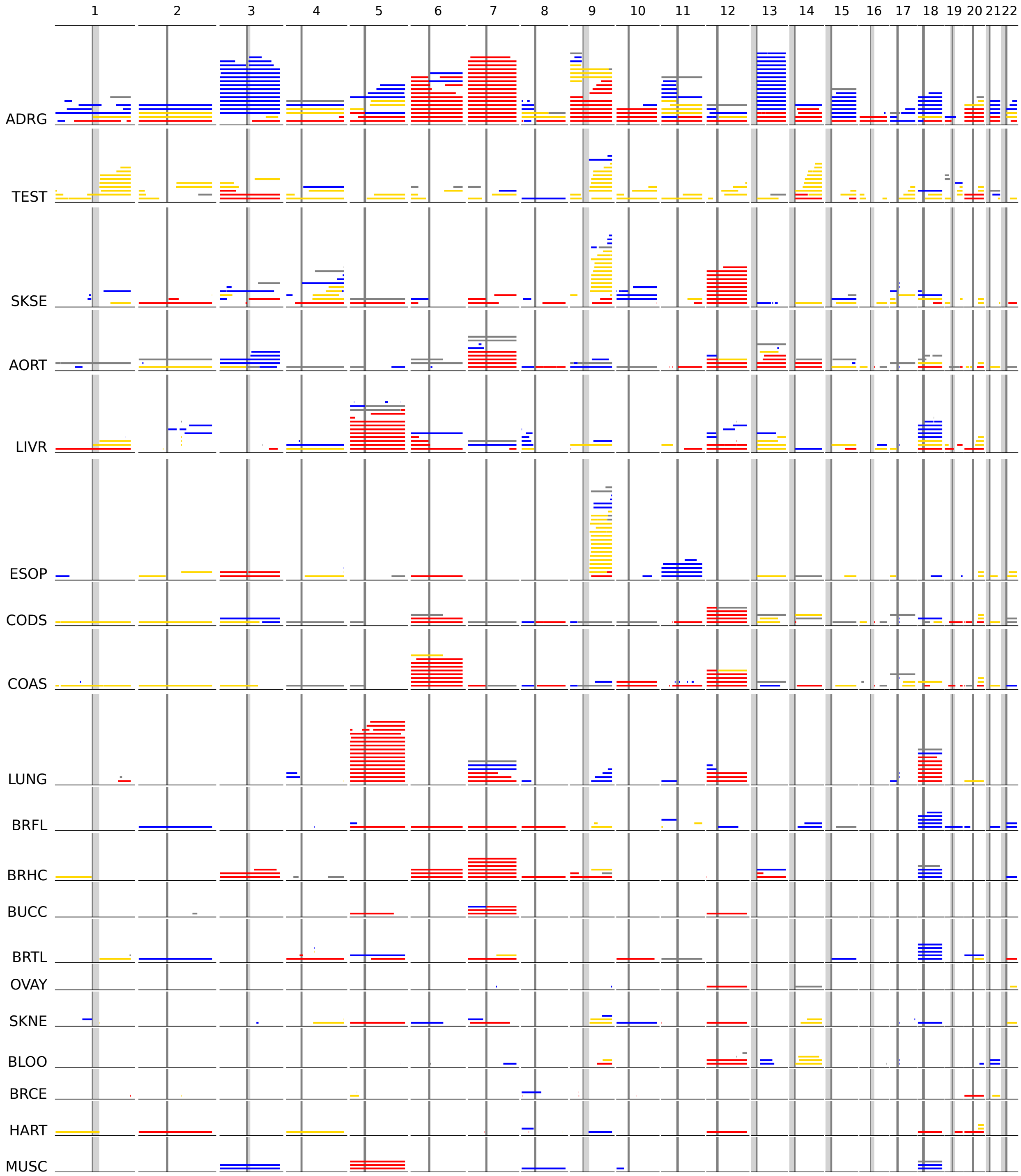


**Supplementary Figure 16: Full view of genomic distribution of autosomal mCAs stratified by tissue**. Each row is a tissue and displays all autosomal mCAs called in that tissue. The columns are the chromosomes and each line depicts the genomic span of an mCA. Within each chromosome, mCAs from the same tissue core share the same y-coordinate. Lines are colored by copy-number state: loss, blue; CN-LOH, yellow; gain, red.

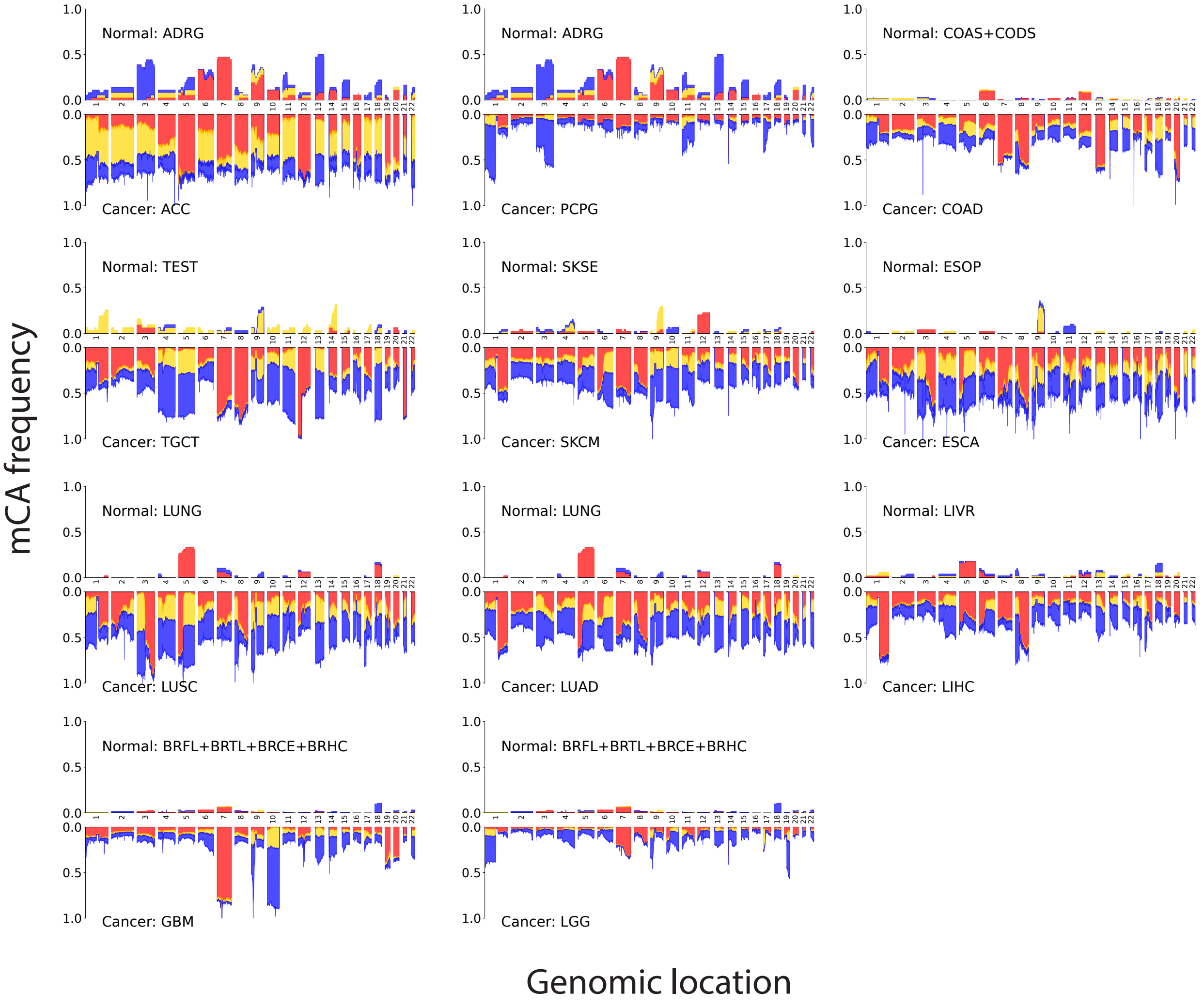


**Supplementary Figure 17: Comparison of mCA landscapes in healthy tissues versus corresponding tumors**. Each panel shows the fraction of samples in SMaHT (top) and TCGA (bottom) with an mCA spanning a given location in the genome. This fraction is broken down into gains (red), losses (blue) and CN-LOH (yellow). SMaHT tissues are paired with cancer types according to the assignment in **Supplementary Table 1**. TCGA data were downloaded from the `files` endpoint of the Genomic Data Commons API (`https://api.gdc.cancer.gov/files`), restricting to 'Allele-specific Copy Number Segment' files produced with ASCAT3 (ref.[4]). For the TCGA data, each segment was assigned an mCA status as follows: CN-LOH if its major copy number was equal to 2 and its minor copy number was equal to 0, loss if its total copy number was less than the median ploidy of the sample, and gain if its total copy number was greater than the median ploidy of the sample. Segments that did not fit into these three categories were left unassigned.

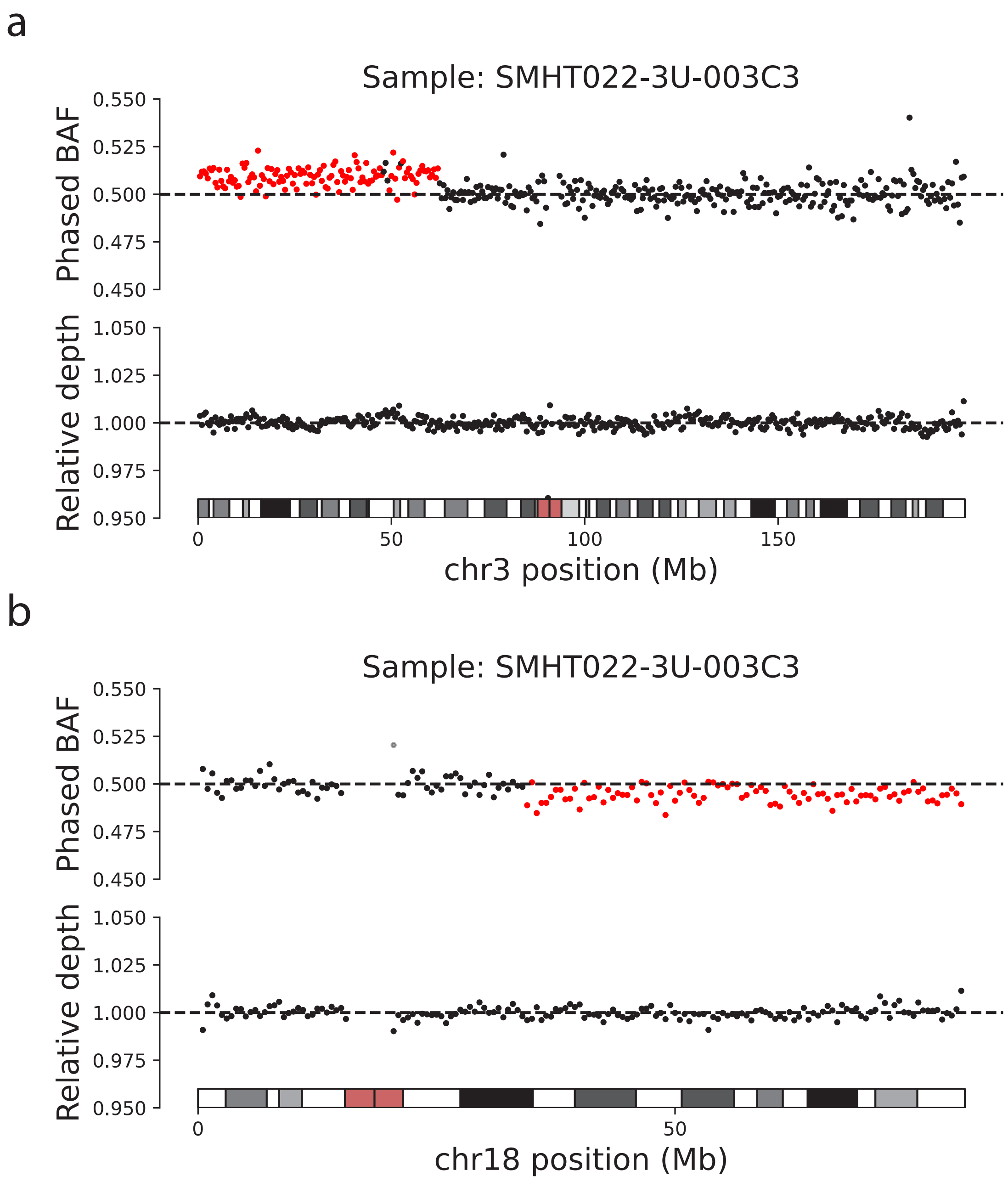


**Supplementary Figure 18: Clean breakpoints of two representative CN-LOH events found in testis**. Measurements of read-depth and phased BAF within each 500-kb window of two chromosomes for a representative testis tissue core are plotted. Dots corresponding to windows within CN-LOH calls are colored red in the phased BAF plots. The step changes in phased BAF are consistent with clonal mutations rather than accumulation of different mutations affecting the same haplotype (which would have different breakpoints, blurring the boundary between allelic balance and imbalance).

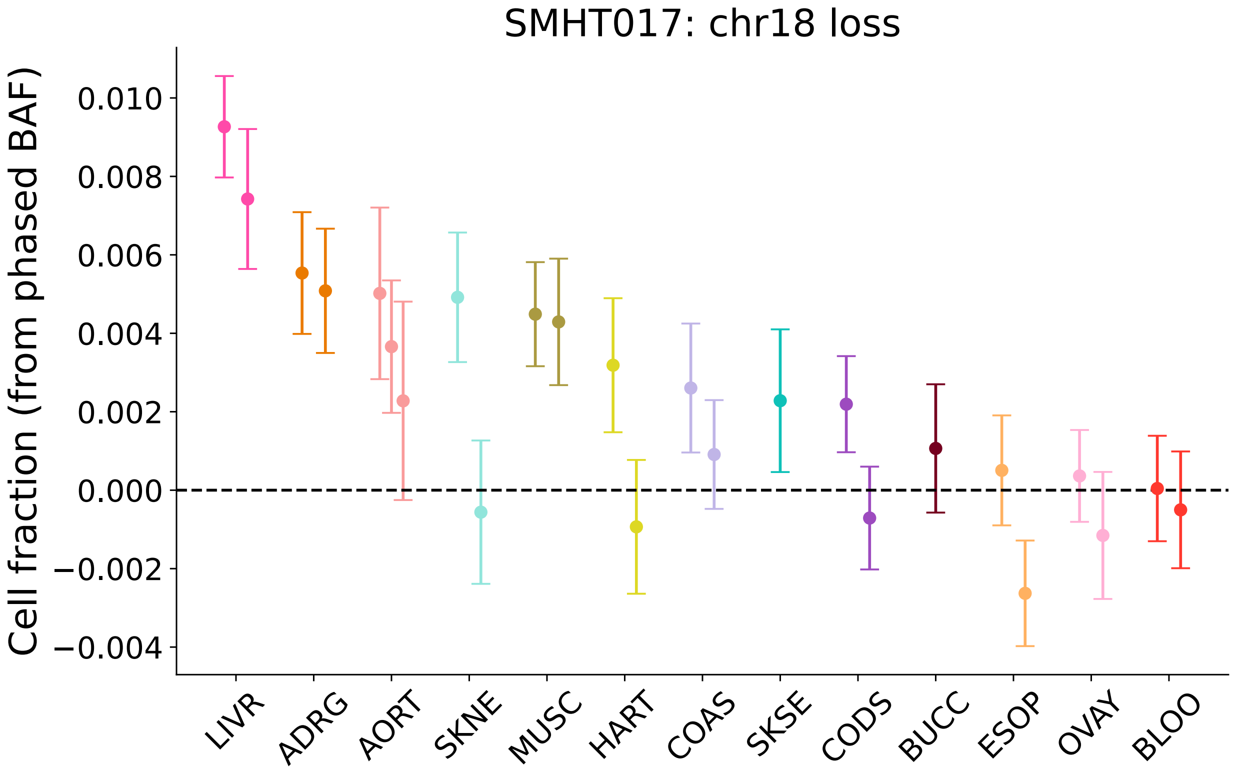


**Supplementary Figure 19: A low cell fraction chr18 loss found in several tissues of SMHT017**. The chr18 loss cell fraction in each tissue is estimated from phased BAF, correcting for phase switches identified in a high cell fraction chr18 gain in lung. Lung is excluded due to the presence of a high cell fraction gain affecting the same haplotype that is lost in other tissues (resulting in an allelic imbalance opposite of that in other tissues).

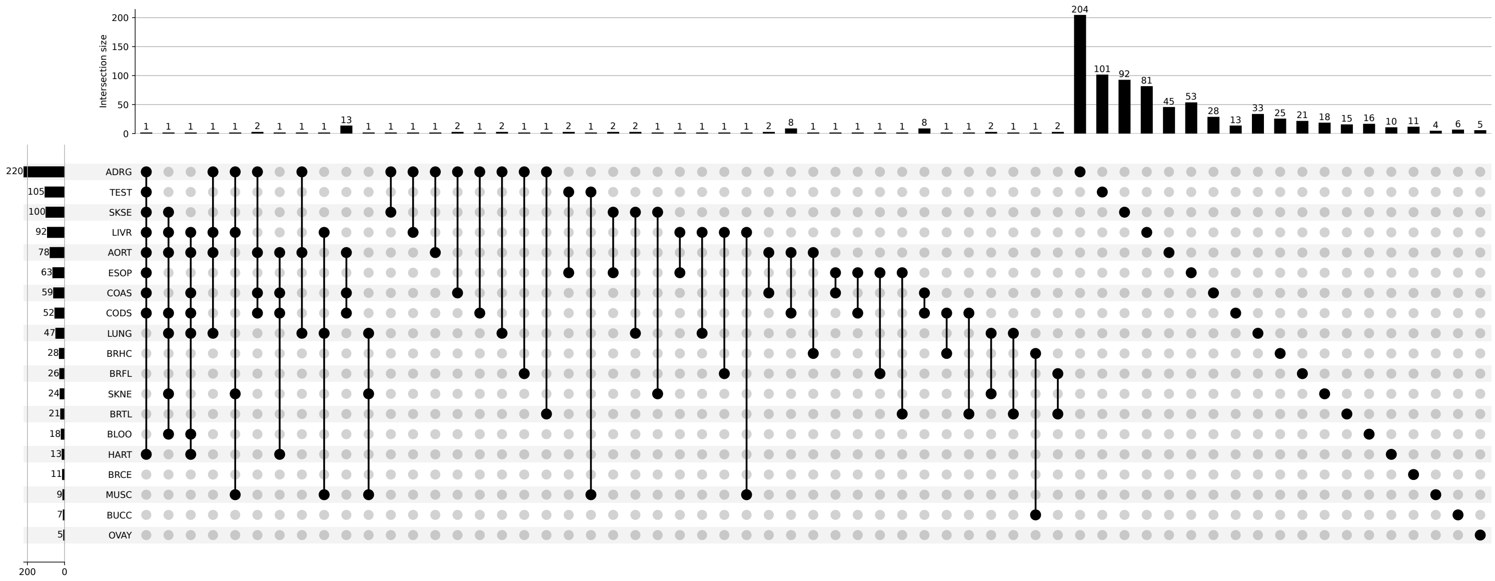

**Supplementary Figure 20: Overview of mCA sharing across tissues within donors**. UpSet plot depicting the frequency of detecting the same mCA in different sets of tissues. Five donors had mCAs shared across 4 or more tissues: the mCAs presented in Fig. 4 and Supplementary Fig. 19, and a clonal population with a complex karyotype that was detected in AORT, COAS, and CODS of SMHT040 (Supplementary Fig. 21). The per row totals do not add up to the total mCAs per tissue since shared mCAs found in multiple cores from the same tissue are only counted once.

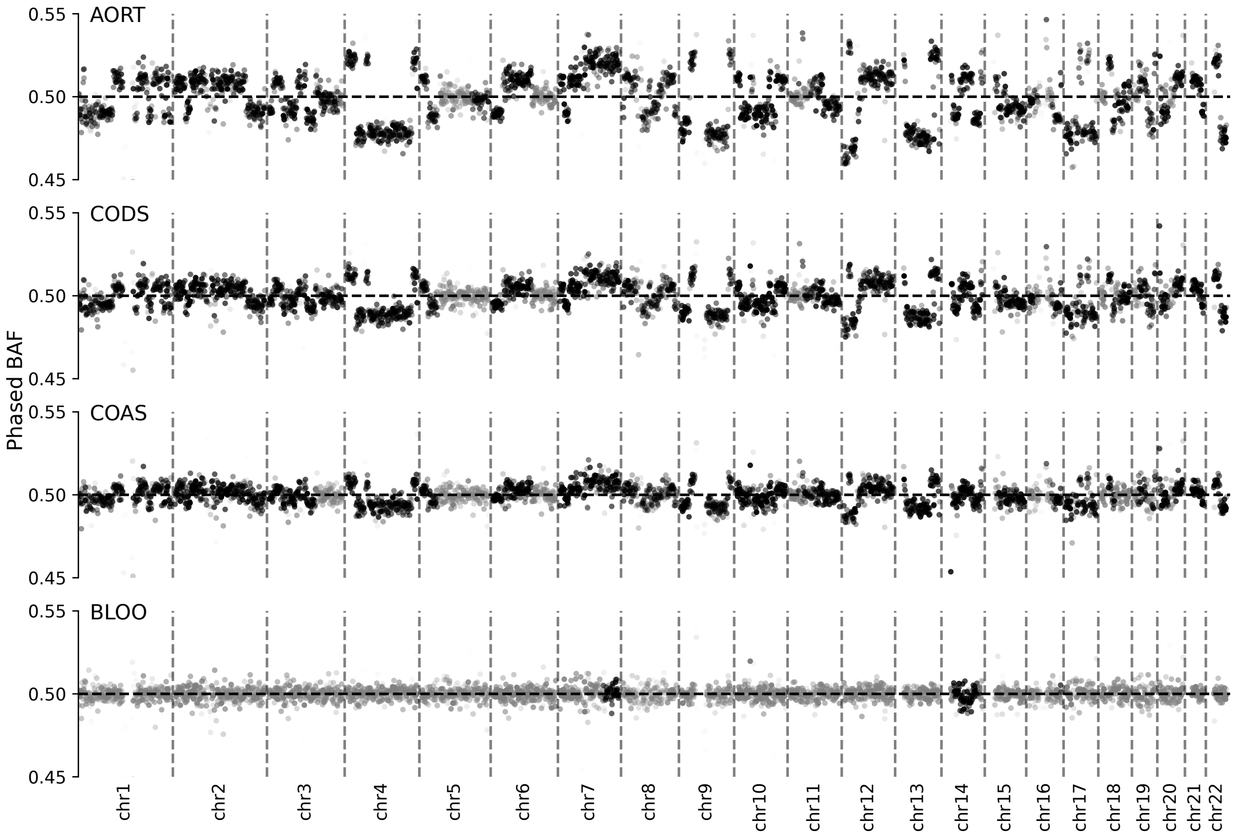


**Supplementary Figure 21: A clone with complex karyotype found in three tissues of a donor**. Genome-wide phased BAF profiles show evidence of a clonal population with complex karyotype in the abdominal aorta and colon (descending and ascending) of SMHT040. Phase agreement among the allelic imbalances in the three cores indicates that the same clone is present in all three samples. The varying levels of allelic imbalance (i.e., absolute deviation from 0.5) within several chromosomes suggest a complex copy number profile. This event was not detected in blood samples from the same donor.

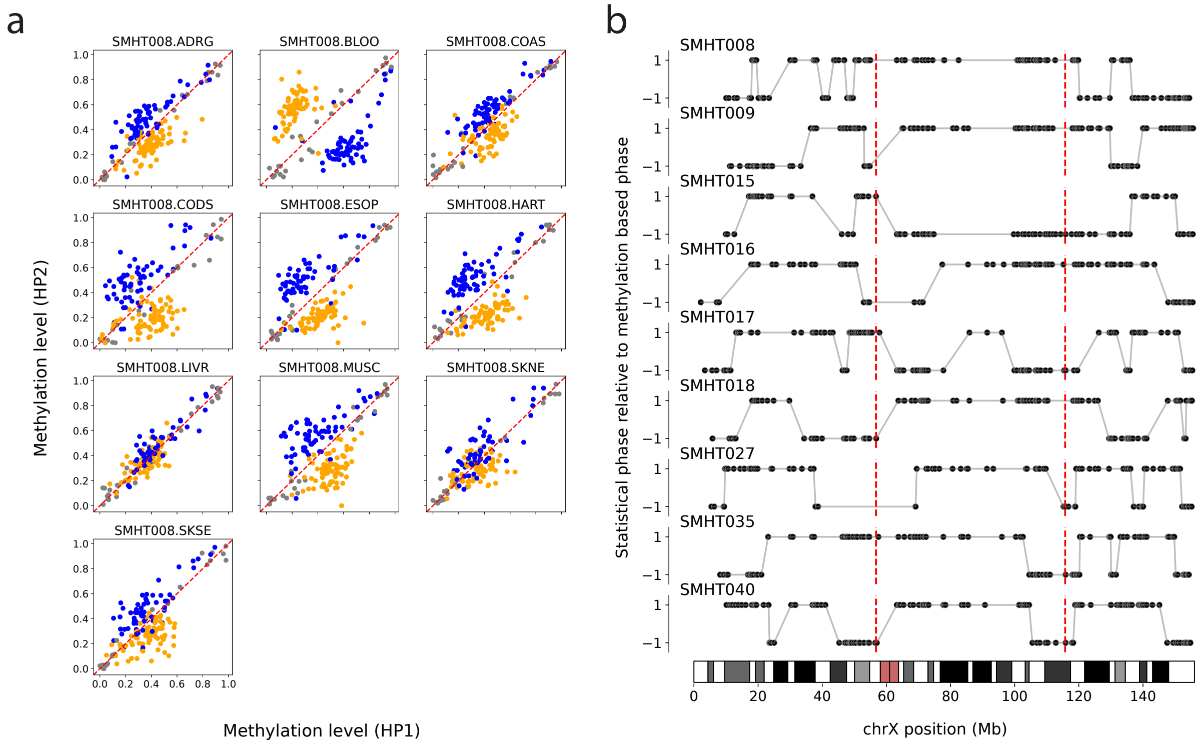


**Supplementary Figure 22: Skewed X chromosome inactivation enables phasing across the X chromosome.** **(a)** Allele-specific methylation data from a representative female donor. Each point corresponds to a long-read-derived phase set; the x- and y-coordinates indicate methylation levels of the two haplotypes (using the initial assignment of haplotypes from statistical plus read-based phasing, which contains switch errors). The symmetric, bimodal distribution of allele-specific methylation levels enables global phasing across the X chromosome since the methylation skew, which reflects skewed XCI, is expected to consistently favor the same homolog across the X chromosome (for a given tissue). A given phase set is plotted in the same color (orange or blue) across all tissues; the color indicates whether its initial assignment of haplotypes was flipped to produce a globally consistent phase. In this example, blood shows the opposite direction of XCI skew compared to the other tissues (see also Supplementary Fig. 23b). Phase sets in gray were not assigned a haplotype and were excluded from subsequent analyses due to lack of a strong methylation skew in any tissue. **(b)** Visualization of switch errors in statistical plus read-based phasing identified using methylation. Each point is an individual phase set plotted at its position along the X chromosome. Each phase set was given a value of 1 or –1 depending on whether or not its assignment of haplotypes was flipped after considering methylation data. Concretely, the orange phase sets in panel **a** were assigned the value 1 while the blue phase sets were assigned the value –1. The red dashed lines indicate the genomic locations of the *ICCE* and *DXZ4* repeat arrays.

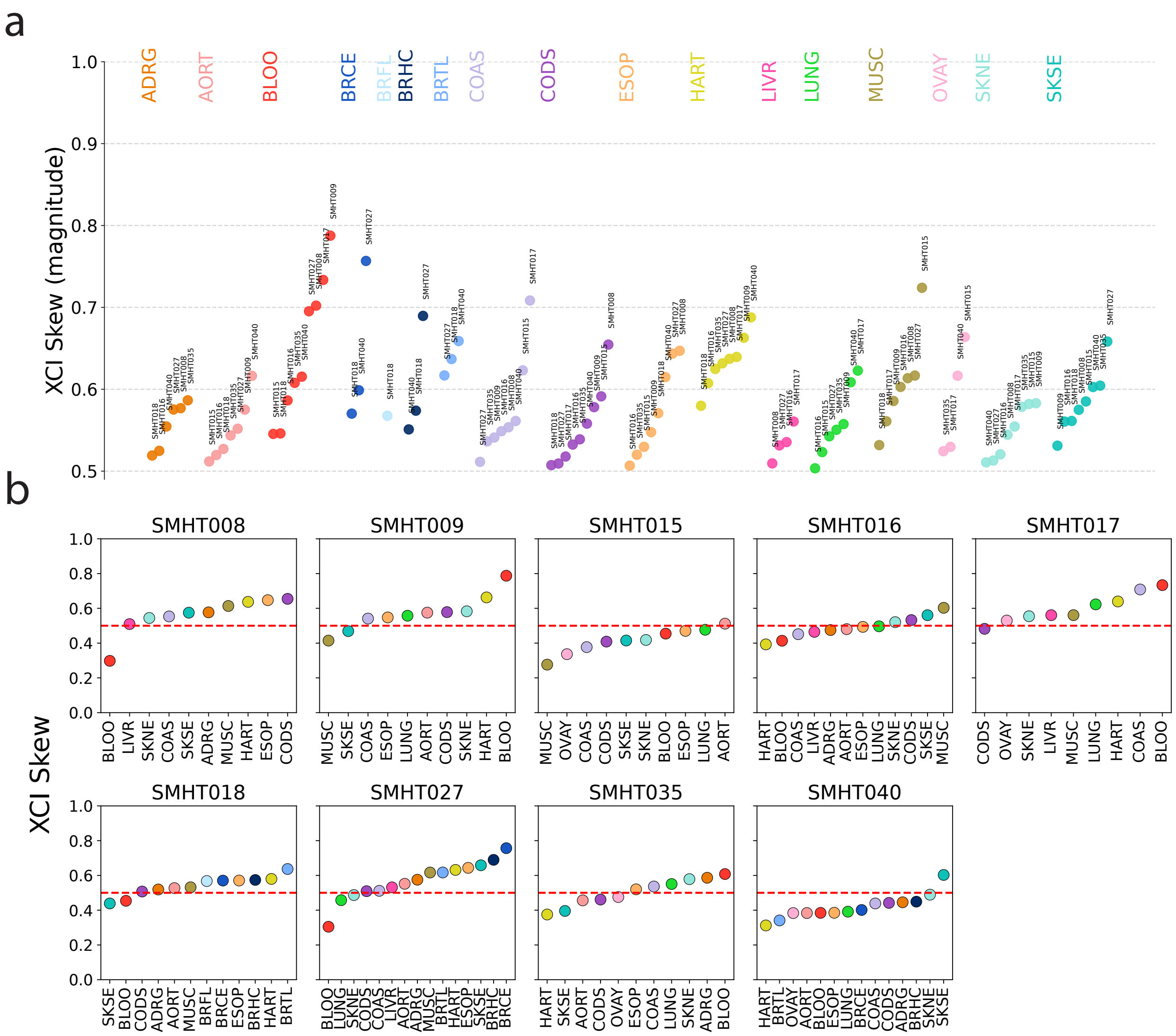


**Supplementary Figure 23: Different tissues from the same donor often display similar X inactivation skew.** **(a)** The magnitude of XCI skew (defined as the fraction of major haplotypes that are inactive) grouped by tissue. All tissues exhibit substantial amounts of XCI skew. **(b)** XCI skews for different tissues from the same donor. XCI skew often favors the same haplotype across most tissues, consistent with early embryonic determination of XCI state.

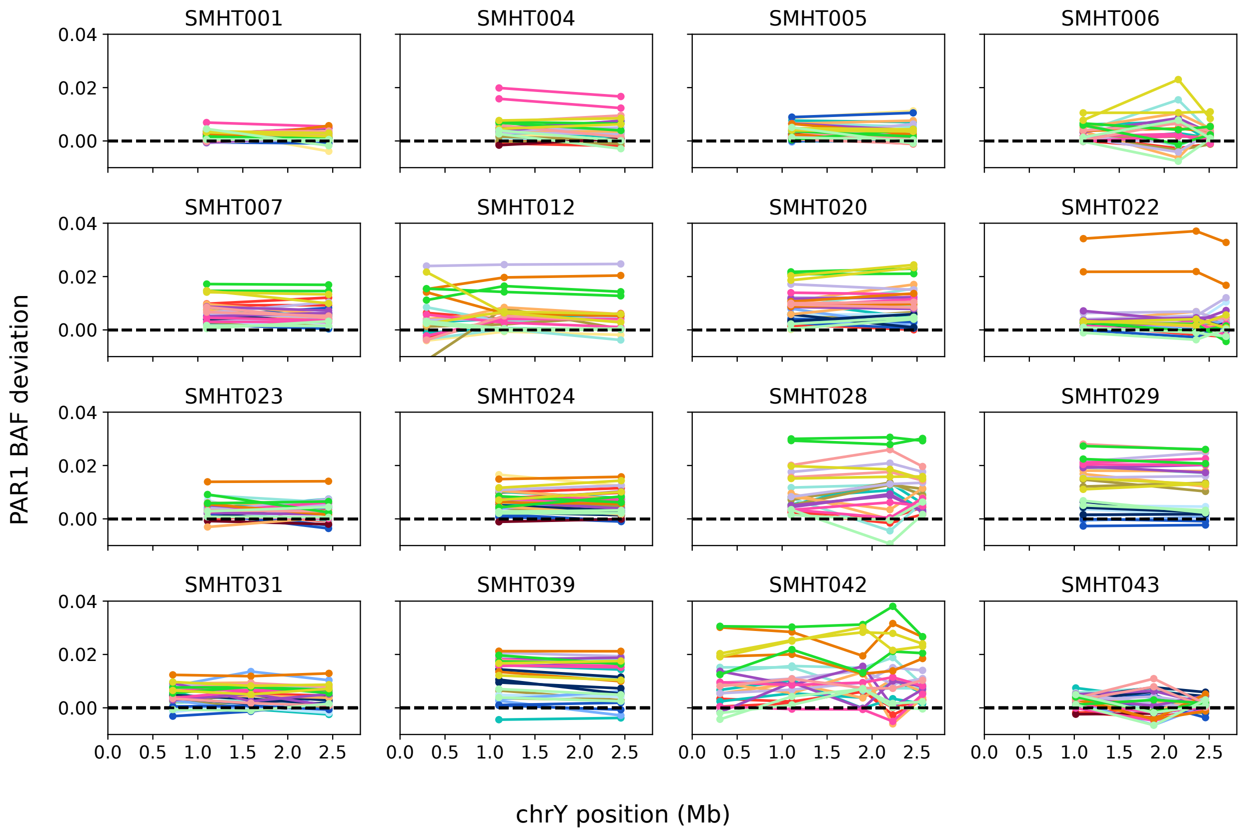


**Supplementary Figure 24: Allelic imbalance is consistent across PAR1 and provides a means to distinguish the Y haplotype from the X haplotype in males**. Each line corresponds to a single tissue core, with each point on the line indicating the allelic imbalance in that core measured using variants within a long-read-derived phase set centered at the position indicated on the x-axis. For most male donors, PAR1 contained only two phase sets, such that each line contains only two points. The global phase for each phase set (i.e., which haplotype corresponds to chrY and which to chrX) was determined by majority vote across tissue cores based on the assumption that the haplotype with lower coverage corresponds to chrY (due to pervasive LOY and essentiality of retaining the X chromosome). Blood samples with large clonal expansions outside the plot range are not shown.

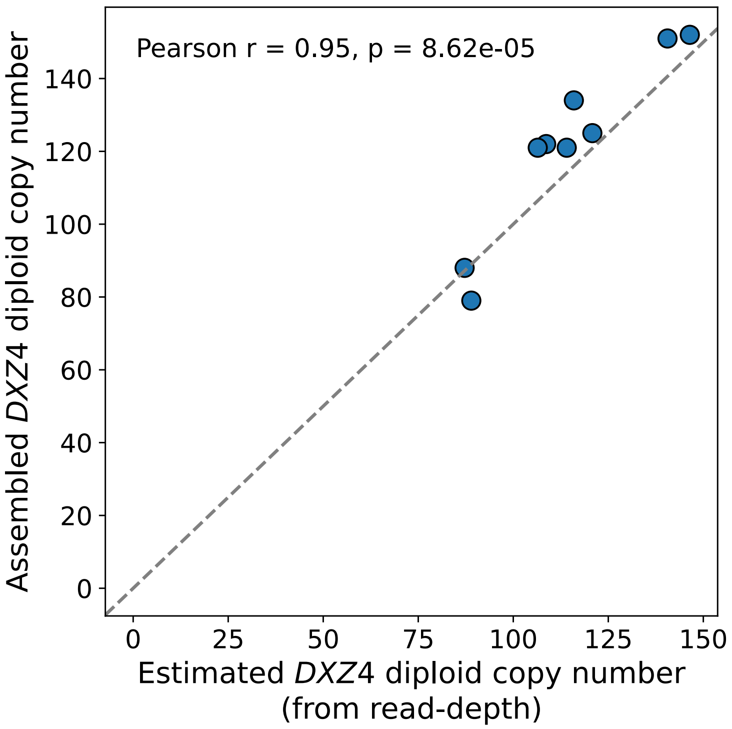


**Supplementary Figure 25: Allele lengths of assembled *DXZ4* alleles agree with estimated diploid copy numbers**. The sum of the two assembled alleles (y-axis) is plotted against the diploid copy number estimated from read-depth in short-read WGS coverage in blood (x-axis) for each female donor.

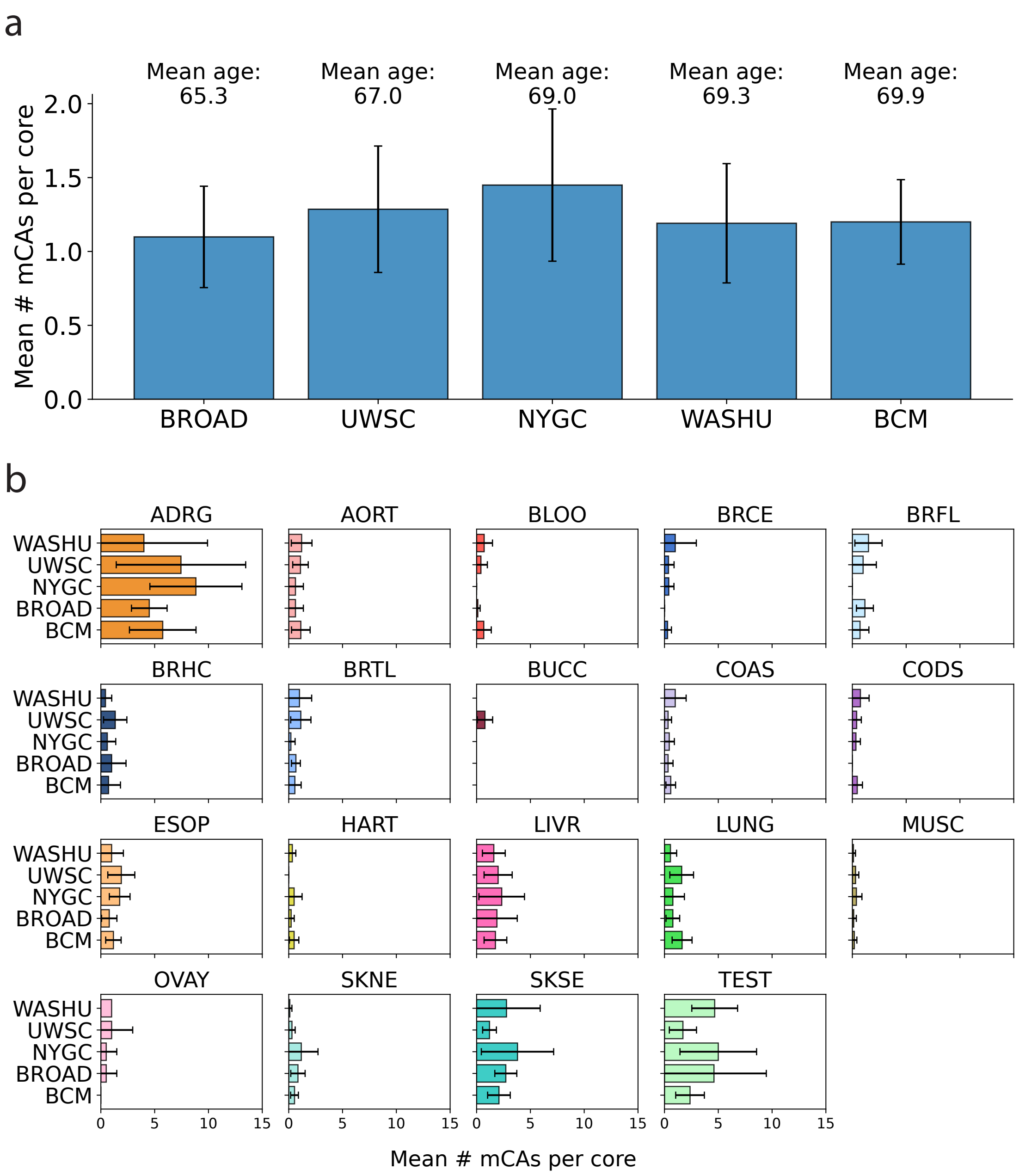


**Supplementary Figure 26: No evidence of sequencing-center-specific batch effects in autosomal mCA calls**. **(a)** Similar rates of mCA detection across tissue cores sequenced at different centers. **(b)** Similar tissue-level mCA burden across sequencing centers. The three outlier cores AORT, COAS, and CODS with the complex-karyotype clone in SMHT040 (sequenced at BCM and UWSC) were removed in both panels. Error bars are 95% confidence intervals estimated as 1.96 times the standard error of the mean. Error bars are missing for OVAY samples sequenced at WASHU due to insufficient sample size (only one core sequenced).
